# DAZL is a master translational regulator of murine spermatogenesis

**DOI:** 10.1101/472191

**Authors:** Haixin Li, Zhuqing Liang, Jian Yang, Dan Wang, Hanben Wang, Mengyi Zhu, Baobao Geng, Eugene Yujun Xu

## Abstract

Expression of *DAZ-like* (*DAZL*) is a hallmark of vertebrate germ cells and essential for embryonic germ cell development and differentiation, yet gametogenic function of *DAZL* has not been fully characterized with most of its in vivo direct targets unknown. We showed that postnatal stage-specific deletion of *Dazl* in mouse germ cells did not affect female fertility, but caused complete male sterility with gradual loss of spermatogonial stem cells (SSCs), meiotic arrest and spermatid arrest respectively. Using the genome-wide HITS-CLIP and mass spectrometry approach, we found that DAZL bound to a large number of testicular mRNA transcripts (at least 3008) at 3′ UnTranslated Region (3′ UTR) and interacted with translation proteins including PABP. In the absence of DAZL, polysome-associated target transcripts, but not their total transcripts were significantly decreased, resulting in drastic reduction of an array of spermatogenic proteins and thus developmental arrest. Thus, DAZL is a master translational regulator essential for spermatogenesis.

---

Germ cells are considered immortal as they are the only cells that pass from one generation to the next, while somatic cells die within a single generation (1). It has long been thought that germ cells utilize a unique set of tools and mechanisms to achieve their distinct function (1, 2). One such tool is conserved germ cell-specific expression of RNA binding proteins among animals (1, 3). Deleted in Azoospermia-like (DAZL) is one of those germ cell-specific RNA binding proteins and its expression is a hallmark of vertebrate germ cells (4-6). DAZL belongs to a human fertility protein family, Deleted in AZoospermia (DAZ) family, consisting of DAZ, DAZL and BOULE (7, 8). DAZ and BOULE appear required only for male fertility (9, 10) but DAZL are required for both male and female fertility (11). Genetic and epigenetic variations affecting human DAZL protein were associated with human infertility (12-15). In addition, DAZL promotes *in vitro* differentiation of human embryonic stem cells towards haploid gametes (16). Hence understanding molecular function of DAZL could provide insight into not only fundamental features of germ cells but also into human infertility and development of the *in vitro* gamete production technology. Mouse Dazl was shown to be critically important in early primordial germ cells (PGCs) and for both male and female fertility (11, 17, 18) in the mixed background of Dazl knockout mice. Analysis of mutant mice in C57/B6 pure background revealed essential roles for development and sexual differentiation of primordial germ cells (PGC) (19-21). Such PGC requirement appear also conserved in other vertebrates (4). By contrast, the role of Dazl in gametogenesis is less clear partly due to the extensive loss of embryonic germ cells in *Dazl* knockout mice. Studies of the remaining spermatogenic cells in *Dazl* mutant testes suggested defects in spermatogonial transition and a final block at leptotene stage of meiosis (17, 18). Stage-specific examination of Dazl function bypassing the early PGC requirement is hence needed to provide a full picture of *Dazl*’s function in gametogenesis.

Dazl has been proposed to be important for mouse oocyte maturation and oocyte-zygotic transition based via *Dazl* RNAi knockdown (22), this is consistent with studies implicating Xenopus Dazl protein’s roles in translational control of oocyte maturation (23-25). Surprisingly, conditional knockout of Dazl by oocyte-specific GDNF9-Cre produced normal amount of pups (26), though the contribution of *Dazl* at or before primordial follicles has not be excluded.

The first clue of DAZ family proteins’ molecular function came from the study on Drosophila *boule* homolog, which was shown to be required for posttranscriptional control of CDC25 homolog, *twine*, (27). While *twine* was not shown to be a direct target of Boule, a series of studies on mammalian DAZL protein revealed potential binding motifs and targets. DAZL bound to the 5’UTR of *Cdc25c* by (GUn)n using SELEX and tri-hybrid system (28). Another *in vitro* experiment (SNAAP) revealed that mDAZL protein could specifically bind to at least 9 distinct mRNAs, including 3′UTR of *Tpx-1* (29). The crystal structures of the RRM from murine DAZL was characterized and verified to be able to bind the motif GUU (30). Reynolds *et al* performed DAZL testis Immunoprecipitation and microarray hybridization and identified at least 11 DAZL targets from the testis directly. Using similar RIP approach, DAZL targets from mouse PGC and human fetal ovaries were also identified (21, 31). However, most of *in vivo* direct targets of DAZL remain unknown. Systematic identification of those direct targets in their physiological context are key to understand how DAZL regulate those targets during spermatogenesis and why DAZ family proteins are critical in sperm development.

While posttranscriptional regulation was established for DAZ proteins, increasing evidence pointed to a major role of DAZL in translational control. Tsui et al. 2000 reported that DAZL could bind to poly(A) RNAs, implicating DAZL in translation control (32). Using a tethering translation assay in *Xenopus laevis* oocytes, Collier *et al* demonstrated nicely that DAZL promote target translation via recruiting PABP (24). Among 11 testicular targets mRNA, Reynolds et al showed that translation of two of them (*Vasa* and *Sycp3*) could be enhanced by DAZL based on a few survived germ cells in *Dazl* knockout mouse testes (33, 34). Polysome profiling of mouse oocytes at different stages revealed extensive translational regulation, DAZL was proposed to be one of the key translational regulators working synergistically with CPEB (22, 35). However translational role of DAZL appear to be context-dependent as both translational promotion and repression were reported for mammalian DAZL (21, 36). Despite the progress on DAZL protein, our understanding of DAZL function in spermatogenesis remained incomplete. A major limitation is a lack of systemic knowledge of its binding targets and interacting proteins in their native context. Comprehensive identification of DAZL targets in gametogenesis hence is necessary to understand their roles and why they are so critical in human fertility. Therefore we determine the stage-specific requirement of *Dazl* in postnatal gametogenesis, then identify comprehensive direct targets and protein partners of DAZL in the mouse testis *in vivo* via both transcriptome-wide HITSCLIP and IP mass spectrometry, and finally interrogate the impact of stage-specific deletion of DAZL on the identified DAZL targets, to unravel the molecular circuitry of DAZL-mediated posttranscriptional regulation.

## Results

### Dynamic and continuous *Dazl* expression from SSCs to round spermatid stage

Although DAZL was found to be highly expressed in various stages of germ cell development using different antibodies and DAZL-GFP reporter previously (8, 11, 37), the extent of DAZL expression, especially in specific stages such as SSC or round spermatids was not clearly defined. We decided to perform a systematic analysis of DAZL expression at both RNA and protein levels using our validated DAZL antibody to gain precise and detailed expression pattern and localization from ES cells through postnatal gametogenesis, supported by Western analysis of Staput-method purified spermatogenic cells of different stages, and immunofluorescent co-localization with stage-specific markers. DAZL protein expression in SSC and round spermatids were clearly established by co-localization with stage-specific markers and western analysis of purified SSC and round spermatids (see Figs Supplementary S1C and 1F). We confirmed DAZL expression in the embryonic gonads and throughout postnatal male germ cell development and differentiation(See Supplementary Fig. S1), validating its expression as a bona fide germ cell marker and suggesting a global and central role during spermatogenesis.

### Removal of *Dazl* in gonocytes, spermatogonia and spermatocyte respectively led to infertility with distinct spermatogenic phenotype

In order to investigate *Dazl* function throughout spermatogenesis, we constructed a conditional *Dazl* knockout mouse and confirmed its nature as a loss of function mutation after deletion of exon 4, 5 and 6, in comparison to the *Dazl* whole body knockout mouse (see Supplementary Fig. S2) (11). The availability of a conditional *Dazl* knockout in combination with germ cell-specific Cre active at different time points of germline development allowed us to determine the requirement of *Dazl* systematically in postnatal gametogenesis. Remarkably, deletion of *Dazl* at gonocyte stage (*Vasa*-Cre; *Dazl*^f/-^, VKO), at spermatogonia stage (*Stra8*-Cre; *Dazl*^f/-^, SKO), and at spermatocyte stage (*Hspa2*-Cre; *Dazl*^f/-^, HKO) (38-40) respectively all led to complete sterility with reduced testis size and absence of sperm in epididymis (see Figs 1A-1D), revealing a persistent critical requirement in at least three stages of spermatogenesis. Examination of adult testis sections showed an absence of germ cells in VKO, an arrest at zygotene stage in SKO, and an arrest at round spermatids in HKO (see Fig. 1E). Hence *Dazl* is critically required in SSC, meiosis and spermiogenesis.

**Figure 1.**
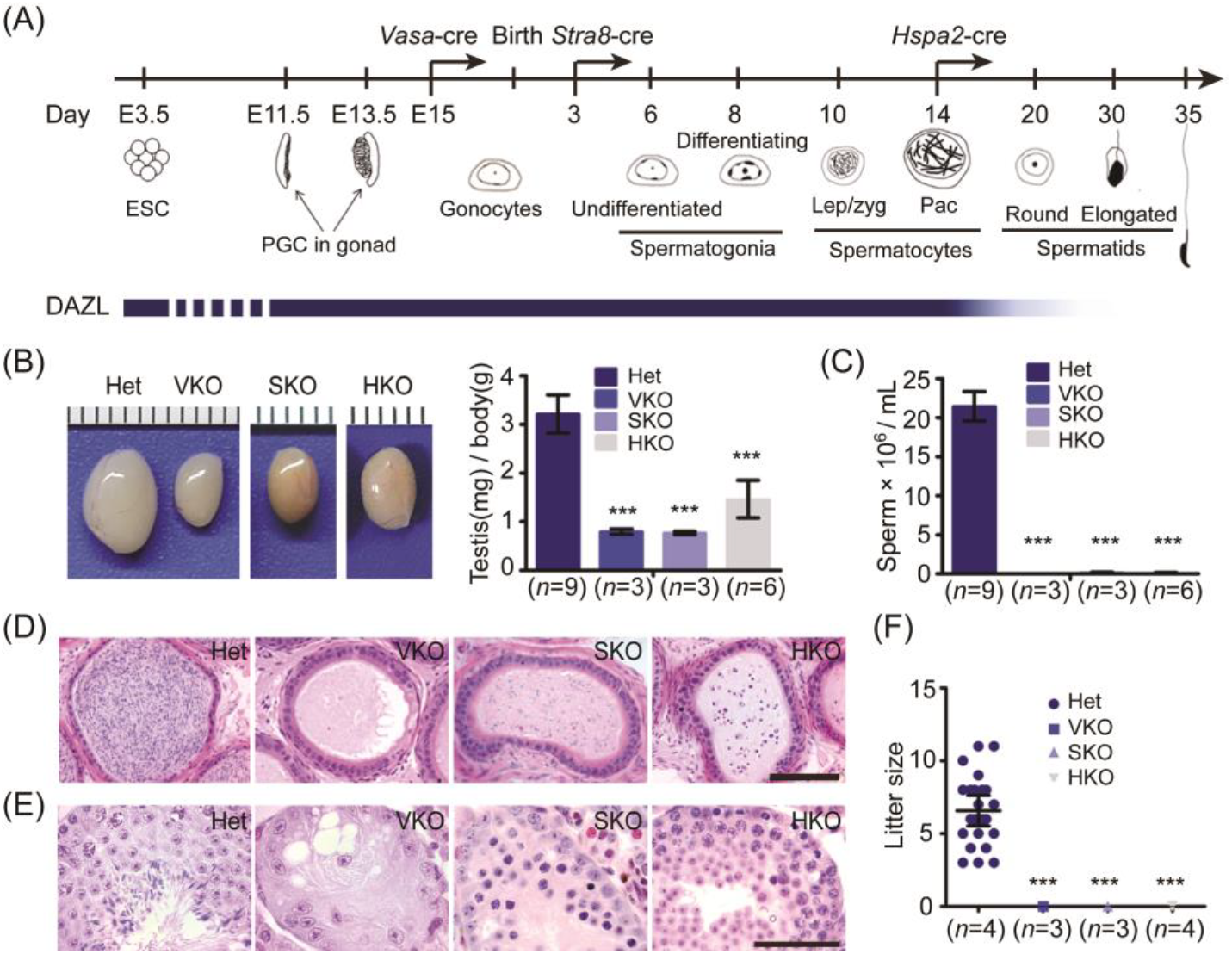
Removal of *Dazl* in gonocytes, spermatogonia and spermatocyte respectively led to infertility with distinct spermatogenic phenotype. (A) Strategy for conditional removal of *Dazl* in gonocytes, spermatogonia and spermatocyte via three germ cell-cre. The onset expression of cre was indicated by an arrow on developmental time line. DAZL protein expression was shown as solid bar in blue color. Dotted line indicated that DAZL expression is unclear during this period. (B) Adult testes size was reduced to different extent in three conditional knockout mice. (C) No sperm in adult VKO and barely any sperm in SKO and HKO were observed in epididymis. (D) H&E stained epididymis sections showed no or little sperm in conditional knockout (cKO) mouse testes. Scale bar: 100μm. (E) H&E stained testis sections from heterozygotes (Het), VKO, SKO and HKO showing distinct spermatogenic defects inside adult seminiferous tubules. Scale bar: 50μm. (F) All *Dazl* cKO mouse were sterile. Error bars indicate SD. *P<0.05, **P<0.01, ***P<0.005.

DAZL is also highly expressed in ovary, in oocytes of all stages (see Supplementary Fig. S3A) (22, 35). Full body *Dazl* knockout led to an ovary without any follicles, similar to the previously reported knockout (see Supplementary Fig. S3B) (11). Surprisingly, both *Dazl* VKO (see Supplementary Figs S3B and S3E, and S3F) and *Gdnf9*-Cre *Dazl*^f/-^ (data not shown) (26) were fertile, with the number of progeny and of pups per litter similar to those of wild type. We confirmed the absence of *Dazl* wild type transcripts and protein in cKO oocytes (see Supplementary Fig. S3C and S3D). To exclude any compensatory effects from *Dazl* paralog, *Boule* (41), we also determined the RNA level of *Boule* transcripts in *Dazl* VKO ovaries. *Boule* transcripts remained undetectable in the absence of *Dazl* (see Supplementary Fig. S3C). Furthermore we constructed *Vasa*-Cre; *Dazl*^f/-^; *Boule*^-/-^ double knockout mouse, those *Dazl* and *Boule* double knockout females remained fertile with no significant difference from wild type or VKO (see Supplementary Figs S3E and S3F). We hence conclude that neither DAZL nor BOULE was required for postnatal female gametogenesis.

### DAZL globally binds mRNAs associated with spermatogenesis at 3′UTR by the motif UGUU

To shed light on how DAZL functions during spermatogenesis, we performed high-throughput sequencing of RNAs isolated by crosslinking immunoprecipitation (HITS-CLIP) to identify DAZL binding targets (42, 43). DAZL antibody was used to pull down RNAs bound by DAZL from a pool of 16 testes from eight 25-day post-partum (dpp) mice. Pulled down RNAs were detected at the predicted DAZL band and at a slightly higher band in the UV cross-linked samples, but not in the non-UV cross-linked samples (see Figs 2A and 2B). The RNAs excised from both the main bands and the slightly higher bands from two independent pull-downs were recovered to construct three cDNA libraries (see Fig. 2C and Supplementary Table S1). The cDNA library constructed from the higher bands overlapped extensively with the cDNA libraries from the predicted main DAZL band, suggesting that the higher bands also contain DAZL targets. Therefore, we used all three libraries for DAZL target analysis. DAZL binding sites appeared to distribute throughout the genome, with the majority of sites (51%) located on the 3′UTR of mRNA, consistent with its role in posttranscriptional regulation (see Fig. 2D). We thus focused our analysis on mRNA targets with DAZL binding sites at the 3′UTR, while intergenic, intronic and noncoding reads will be analyzed separately. And predominant binding sites for mRNAs are located on 3′UTR. Out of 1373 shared mRNA targets among all three libraries, 1235 targets contained binding sites at the 3′UTR (see Supplementary Figs S4A and 2E). For mRNAs shared by at least two libraries, 3008 out of 3470 targets contained binding sites at the 3′UTR. Those targets (1235 or 3008 targets) were significantly enriched for pathways involved in RNA metabolism, such as mRNA processing, RNA splicing and translational regulation. Cell cycle and spermatogenesis pathways were also highly enriched (see Fig. 2F). To determine the validity of the DAZL target mRNAs identified by HITS-CLIP, we randomly picked 8 targets from among the 1235 3′UTR targets shared by all three libraries and 7 targets from among the less stringent 3008 3′UTR targets (shared by at least two libraries) and found they were all significantly enriched in DAZL immunoprecipitation (IP) (see Fig. 2G). By contrast, Sertoli cell transcripts *Wt1*, *Gata4* and other non-targets such as *Nanos2*, *Neurog3* (*Ngn3*), *Tnp1* and *Tnp2* were not enriched (see Figs 2G and Supplementary Fig. S4B). This suggested that even the 3008 shared targets were reliable DAZL targets. We further established the quality and reproducibility of the three libraries by comparing the binding sites of the targets. We observed a consistent peak distribution for the same target transcripts among the three libraries (see Supplementary Fig. S4C). To identify the consensus sequence of DAZL binding sites, we used a motif discovery algorithm HOMER (44) to search for the mRNA motifs bound by DAZL. Of the most enriched 3-mer motifs, the GUU triplet ranked on the top in all three samples with much higher statistical significance than other motifs, consistent with the binding motif determined by crystal structure and binding affinity analysis of DAZL RRM-RNA complex (30) (see Fig. 2H). Among the most enriched 6-mer motif, the VVUGUU ranked on the top. To test the UGUU binding motif, we evaluated the 3′UTR of the DAZL target *Sycp1* using a dual luciferase assay (see Fig. 2I). When we mutated UGUU to ACAA, relative luciferase activity decreased significantly, confirming the consensus binding motif of DAZL (see Fig. 2J).

**Figure 2.**
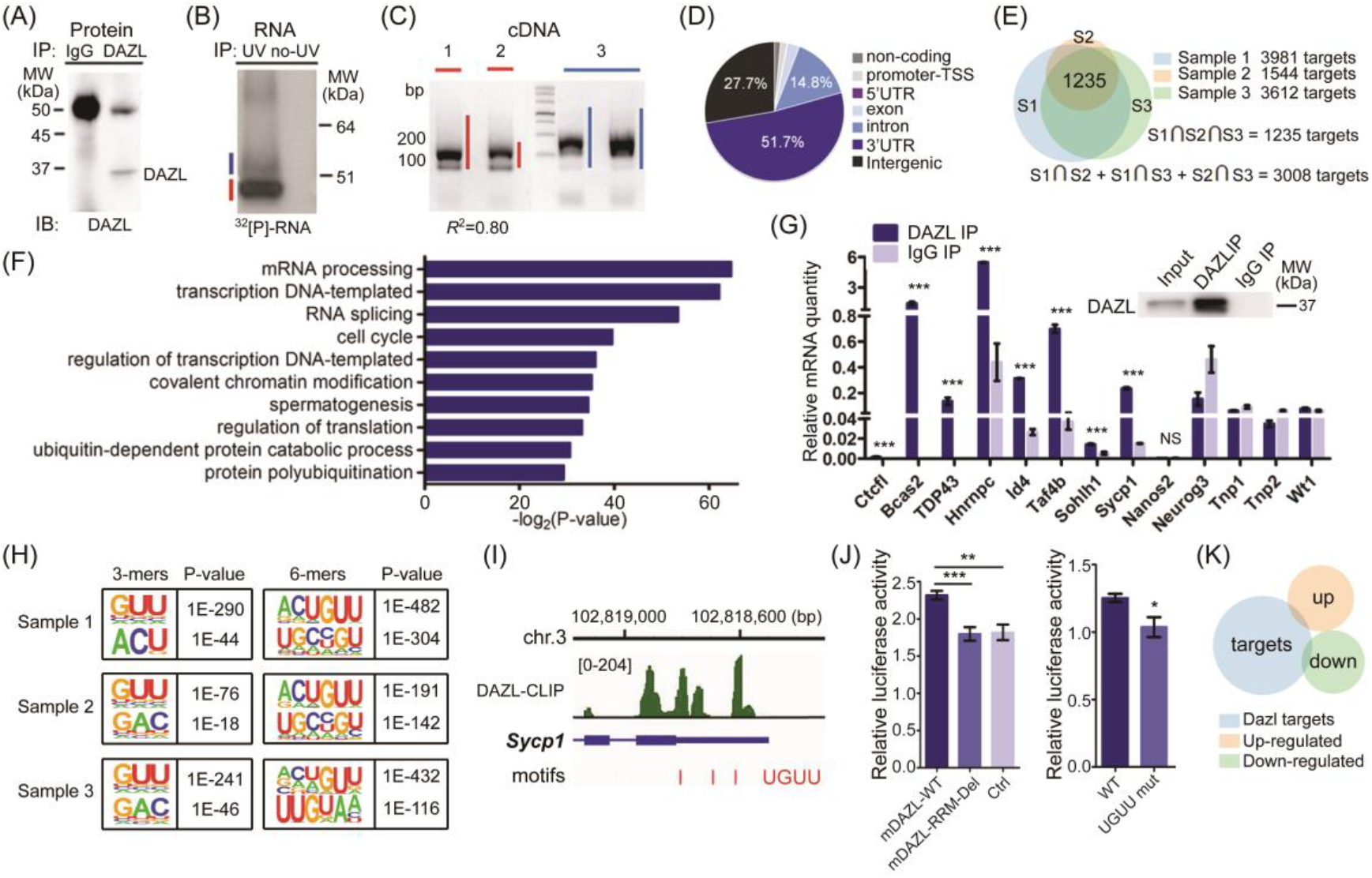
Identification of DAZL targets in mouse testes by HITS-CLIP. (A) Immunoblot (IB) analysis of DAZL immunoprecipitation (IP). (B) Autoradiogram of ^32^P-labeled RNA crosslinked to DAZL pulled down by IP. The RNA marked by blue and red bars was excised for library construction. (C) RNA excised from (B) was ligated to linker (36nt) and amplified by PCR. The amplified cDNA product marked by blue and red bars was derived from the band of the same color in (B). (D) Distribution of all DAZL binding regions based on their genomic locations. (E) Venn diagram of DAZL targets bound at 3′UTR from three cDNA libraries, including two main bands (sample 1 and 2) and a mixture of mRNA combined from the two higher bands (sample 3). (F) Gene ontology analysis result of DAZL targets revealed by DAVID(Huang da, 2009 #9303;Huang da, 2009 #5426). (G) Validation of HITS-CLIP result by RNA Immunoprecipitation (49) of 8 target genes. (H) Consensus motifs within DAZL clusters from 3 samples identified by HITS-CLIP using the HOMER algorithm. (I) Consensus motif distribution at the 3′UTR of a DAZL target *Sycp1*. (J) DAZL bound to the 3′UTR of *Sycp1* around the motif ‘UGUU’ and enhanced luciferase expression. Error bars indicate SD. *P<0.05, **P<0.01, ***P<0.005. (K) DAZL target genes showed little overlap with up or down regulated genes in 10dpp *Dazl* SKO testes.

### DAZL does not significantly impact the stability of its targets mRNA

We next examined transcript abundance of the identified DAZL targets in the absence of DAZL. To our surprise, there was little overlap between the DAZL targets identified by HITS-CLIP (see Fig. 2K) and genes whose transcript level significantly changed in the 10dpp *Dazl* SKO testes. We chose 10dpp testes because knockout testes were comparable to those of control in cell composition. And lack of transcript level change in the knockout testes among DAZL targets could not be attributed to the different time points of testes used, as majority (2896 out of 3008) of DAZL targets were detectable (FPKM >30) at 10dpp testes. This result suggested that loss of DAZL did not significantly impact target transcript levels, leading us to propose that mouse DAZL regulates translation rather than transcript stability. Furthermore, changes in transcript levels of non-target genes in *Dazl* mutant testes compared to WT testes were likely a secondary effect of *Dazl* loss.

### Protein expression of DAZL SSC-associated targets is essential for maintenance of spermatogonial stem cells

Since spermatogenic genes were highly enriched among DAZL targets, we mapped all the DAZL binding sites of spermatogenic targets. Remarkably, all the spermatogenic target mRNAs contained binding sites exclusively on their 3′UTR (124 out of 124, classified by DAVID, see Supplementary Table S2), consistent with DAZL being a translational regulator (45). We also found that DAZL target transcripts were present throughout spermatogenesis, with enrichment for genes associated with key steps such as SSC self-renewal, meiosis and spermatid differentiation. Such stage enrichment corresponded nicely with the stages *Dazl* shown to be required from our genetic analysis above, leading us to hypothesize that DAZL may promote spermatogenesis by regulating protein expression of many target transcripts critical for each stage of spermatogenesis.

Among the spermatogenic mRNAs bound by DAZL, there was a remarkable enrichment of genes linked to SSCs. Out of 13 genes shown to be important for maintenance of the spermatogonial progenitor pool (46), the mRNAs for 8 of these genes were found to be DAZL targets (see Fig. 3A), suggesting a previously unknown function of DAZL in the regulation of SSCs. Careful examination of binding peaks among those 8 genes revealed distinct peaks on their 3′UTRs (see Fig. 3B). We then determined DAZL expression in SSCs and found that both DAZL mRNA and protein were detectable in neonatal testes and in purified primary SSCs, and DAZL co-expressed with PLZF^+^ and LIN28A^+^ cells (see Supplementary Figs S1A-C and F), further supporting a role of DAZL in SSCs (47).

**Figure 3.**
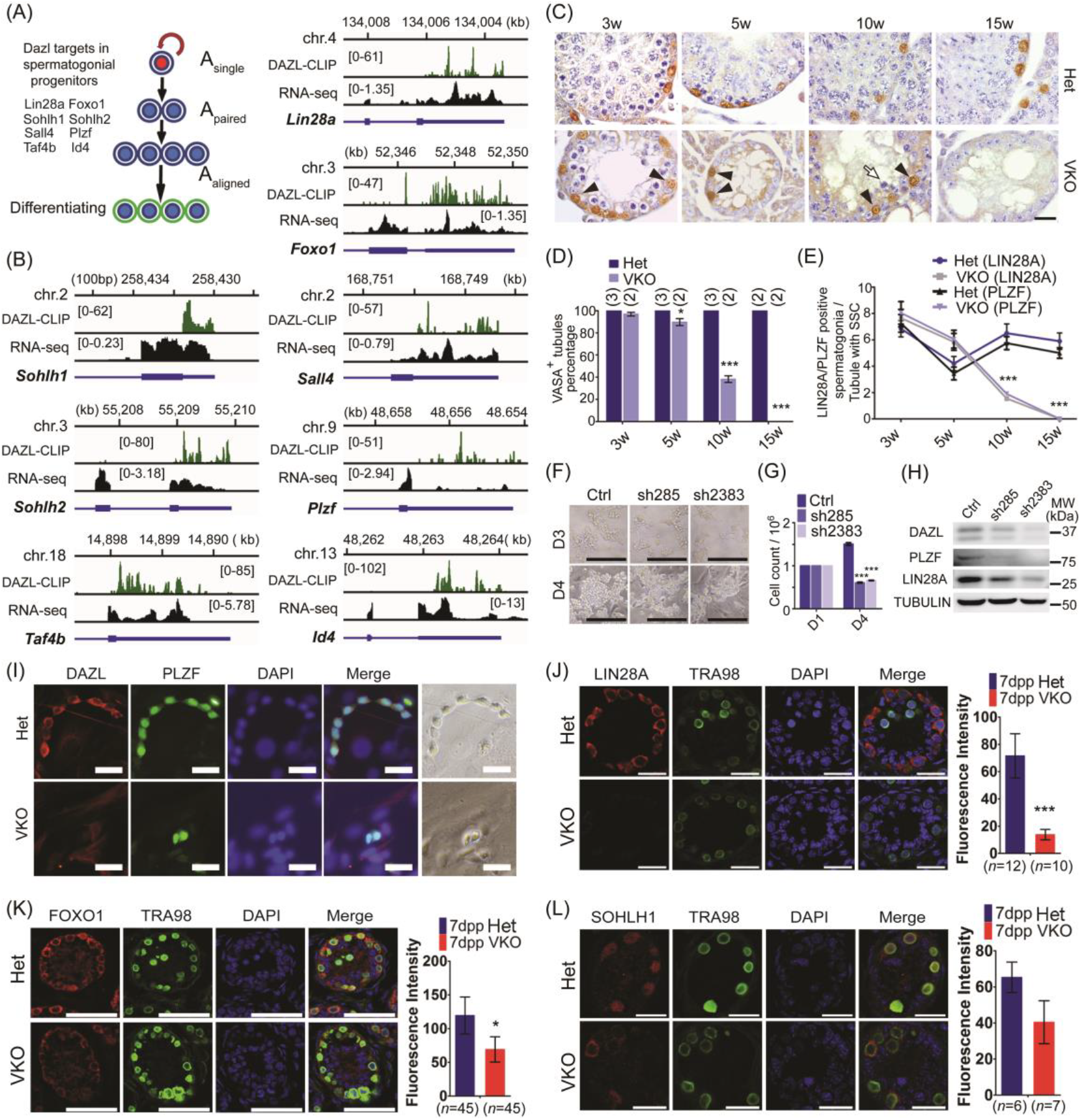
DAZL targets are enriched for SSC-associated genes and loss of DAZL leads to a defect in spermatogonial progenitor maintenance. (A) Schematic illustration of SSC-associated genes bound by DAZL. (B) Genome browser tracks showing binding peak distributions (DAZL-CLIP) and transcript levels (RNA-seq) of SSC-associated target genes. (C) Immunohistochemical staining of the SSC marker PLZF in *Dazl* VKO testes. Scale bar: 20 μm. (D) Gradual germ cell loss in testes of *Dazl* VKO mice. Testes sections from 3, 5, 10 and 15-week-old mice were stained for VASA expression to detect the number of VASA-positive tubules. A seminiferous tubule containing at least one VASA-positive cell was classified as a VASA-positive tubule. The number of mice at each time point is shown in parentheses. More than 100 cross-sections were scored for samples taken from random slides. (E) Comparison of the number of LIN28A^+^ and PLZF^+^ SSCs in cross-sections of testes from *Dazl* heterozygotes (Het) and *Dazl* VKO mice at different ages. The number of mice is the same as in (D). A seminiferous tubule containing at least one SSC was classified as tubule with SSCs. Data are mean ± SEM for 20 round seminiferous tubules of both genotypes at each age. *P<0.05, **P<0.01, ***P<0.005. (F) *Dazl* knockdown led to a significant reduction in the size of SSC clones at day 4 following shRNA lentiviral transduction. Scale bar: 200 μm. (G) Cell count of SSCs at day 4 following lentiviral transduction. (H) SSC markers (LIN28A, PLZF) were downregulated at day 4 following lentiviral transduction. (I) Established SSC clones immunostained for DAZL and PLZF after 4 days of culture of Thy1^+^ cells from 7dpp *Dazl* heterozygotes and *Dazl* VKO testes. Scale bar: 50 μm. (J-L) Fluorescence intensity of DAZL targets were compared between *Dazl* heterozygotes and *Dazl* VKO testes at 7dpp by immunofluorescent staining in TRA98-positive cells. Scale bar: 20 μm. Error bars indicate SEM. *P<0.05, **P<0.01, ***P<0.005.

We then investigated the effect of removing *Dazl* from SSCs by generating *Dazl* VKO mice; these mice were expected to produce no functional DAZL in postnatal germ cells, including SSCs (38). In testes from 3-week-old *Dazl* VKO mice, despite a significant reduction in germ cells, the number of both LIN28A^+^ and PLZF^+^ cells were comparable to controls, suggesting that the initial pool of SSCs was not different between heterozygotes and knockout mice. By 5 and 10 weeks of age, however, LIN28A^+^ and PLZF^+^ cells had decreased gradually and completely disappeared by 15 weeks of age (see Figs 3C-E, Supplementary Fig. S5A, and S5B), resembling the reported phenotypes of SSC maintenance-defective mutant mice (48). shRNA knockdown of *Dazl* in established SSC cultures led to significantly fewer SSC clones with reduced size and cell number (see Figs 3F and 3G). PLZF and LIN28A proteins were also significantly reduced in the *Dazl* knocked-down SSCs, establishing a role of DAZL in SSC self-renewal and development, by promoting protein expression of SSC-associated target genes (see Fig. 3H).

To examine the self-renewal ability of *Dazl* knockout SSCs, we isolated THY1^+^ cells from *Dazl* VKO testes to generate *in vitro* SSC clones without DAZL protein. The *Dazl* knockout SSC clones were established *in vitro* and had similar morphology as heterozygotes SSCs in the first and second day of culture (data not shown). However, the knockout SSC clones failed to grow further, forming only chains or clusters of 2-3 cells; by contrast, the heterozygotes SSCs formed long chains and large clusters (see Figs 3I and Supplementary Fig. S5C). These data demonstrated an essential role of DAZL in the maintenance of the spermatogonial progenitor pool.

Furthermore, we examined protein expression of DAZL targets in *Dazl* knockout germ cells. Based on fluorescence intensity relative to germ cell marker TRA98, the expression of SSC-associated DAZL target proteins was decreased in 7dpp *Dazl* VKO mouse testes, with LIN28A and FOXO1 showing a statistically significant decrease compared to controls (see Figs 3J-L). This finding further supports that DAZL-mediated SSC-associated protein expression is critical for SSC maintenance.

### DAZL-mediated protein expression of meiotic genes is essential for synaptonemal complex (SC) assembly and DNA repair during meiosis

Other spermatogenic genes significantly enriched in the DAZL targets were associated with meiotic cell cycle (see Figs 4A and 4B). The list of DAZL-targeted meiosis-associated genes included the previously reported *Sycp3* as a direct target (34), but contain a number of other meiotic genes such as major synaptonemal complex genes (*Sycp1* and *Sycp2*) and DNA repair pathway genes (*Rad51, Msh4*,etc). We investigated the effect of loss of *Dazl* on the expression of these DAZL targets, as well as on key meiotic events. To bypass the embryonic and SSC reqiurement for *Dazl*, we utilized *Stra8*-Cre, which drives Cre expression starting in the 3dpp testis (39). We found that the resulting *Dazl* SKO mice are sterile and have significantly smaller testes compared to heterozygotes (see Figs 1B and 1C). Spermatogenesis was arrested at the transition from zygotene to pachytene stage in adult testes, and in the first wave of spermatogenesis of both *Dazl* VKO and *Dazl* SKO animals (see Supplementary Figs S5A and S6A), confirming a meiotic block in the absence of *Dazl* (18). Expression of the SSC marker PLZF and early meiotic markers such as STRA8 and SPO11 were not affected in 9dpp *Dazl* SKO mice, suggesting that those knockout germ cells were able to enter meiosis (see Fig. 4C). Indeed, meiotic chromosome spread revealed a similar proportion of leptotene cells in the *Dazl* SKO mice compared to wild type but significantly more zygotene and pachytene-like cells, suggesting an arrest at the transition from zygotene to pachytene (see Supplementary Fig. S6B).

**Figure 4.**
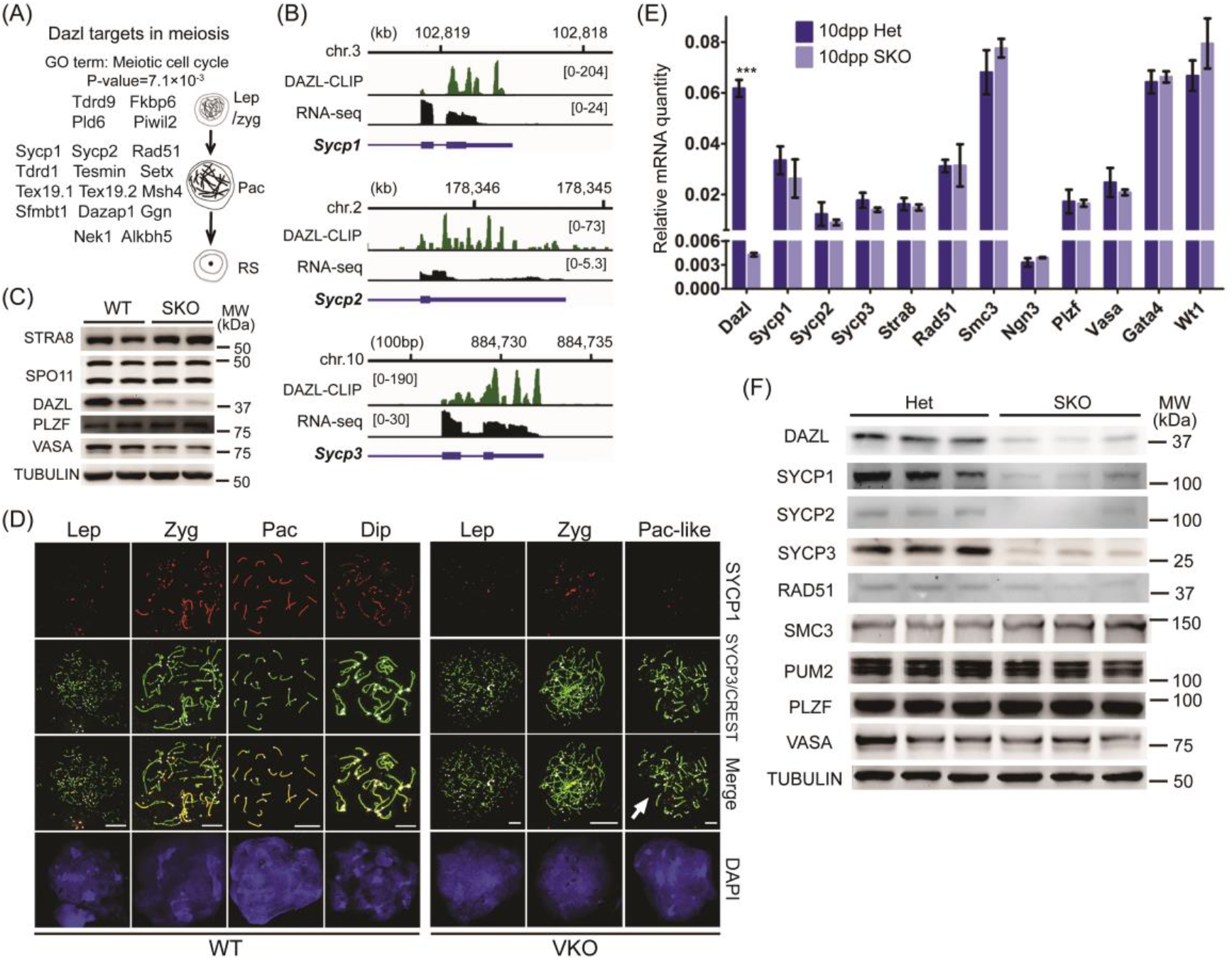
Synapsis defect in *Dazl* knockout spermatocytes results from reduced protein expression of DAZL targets *Sycp1, Sycp2* and *Sycp3*. (A) Schematic illustration of meiotic genes bound by DAZL. (B) Genome browser tracks showing binding peak distributions and transcript levels of meiotic target genes of the synaptonemal complex. (C) Expression of proteins involved in meiotic initiation and SSC-associated proteins were not significantly changed in 9dpp *Dazl* SKO testes. (D) Meiotic spread of 21dpp wild type and *Dazl* VKO spermatocytes co-stained with SYCP1, SYCP3 and CREST antibodies. Completion of synapsis is determined by a single CREST signal (white points) for a given pair of chromosomes. (E) Transcript levels of meiotic target genes were not changed significantly in 10dpp *Dazl* SKO testes. (F) Protein level of meiotic target genes were significantly reduced in 10dpp *Dazl* SKO testes.

In DAZL-deficient leptotene nuclei, short stretches of condensed SYCP3 signal were observed, indicating that chromosome core-associated SYCP3 protein was loaded normally during early meiotic prophase. However, loss of *Dazl* resulted in an increase in the unpaired synaptonemal complex in zygotene stage spermatocytes (see Fig. 4D). Despite the presence of SYCP3-positive axial elements, nearly all of the *Dazl* VKO zygotene spermatocyts had weak SYCP1 staining, suggesting a partial loss of transverse filaments of the synaptonemal complex (white arrow, see Fig. 4D). Defects in synapsis formation were also observed in *Dazl* VKO pachytene spermatocytes. Typical pachytene nuclei with 20 synapsed chromosomes were never observed in *Dazl* VKO cells. Instead, most nuclei contained unsynapsed 40 chromosomes (n=222) with fewer than 5% exhibiting partial synapsis. The vast majority of axial elements in *Dazl* VKO spermatocytes failed to align and instead remained separated from each other as univalent chromosomes. The presence of about 40 short SC suggested that the chromosomes were condensed and that the spermatocytes were at a stage corresponding to early pachytene; we refer to these as ‘pachytene-like.’ The same abnormal pachytene-like cells were found in *Dazl* SKO testes (see Supplementary Fig. S6C). These results suggest that the loss of *Dazl* disrupted the assembly of the synaptonemal complex, and likely led to meiotic arrest at early pachytene.

In addition to defective synapsis and disruption of the synaptonemal complex, we also observed increased DNA damage in *Dazl* VKO spermatocytes. Phosphorylated H2AX (called γ-H2AX or H2AFX) is a marker of double strand breaks (DSB) during leptotene and zygotene, and of the XY body during pachytene (49, 50). While *Dazl* VKO leptotene spermatocytes showed similar staining for H2AFX, the signal did not decrease in zygotene, and instead increased in both zygotene and pachytene-like cells. The single XY body observed in wild type pachytene spermatocytes was not present in the *Dazl* VKO pachytene spermatocytes. A number of dispersed H2AFX domains were retained along each chromosome (see Supplementary Fig. S6D). A similar dispersed H2AFX staining pattern was seen in *Dazl* SKO spermatocytes, implicating DNA repair defects in the absence of *Dazl* (see Supplementary Fig. S6C). DAZL meiotic target mRNA expression was not affected in the SKO testes but their protein level was significantly reduced (see Figs 2K and 4E, and 4F), consistent with a defect in the synaptonemal complex, synapsis and DNA repair in the mutant testes. These findings further support a role for DAZL in translational regulation rather than mRNA stability in the regulation of spermatogenesis.

### DAZL is required for translation of its targets mRNAs and interacts with PABP in mouse testes

Given the requirement of DAZL throughout spermatogenesis, we next examined the mechanism of DAZL action at the molecular level, specifically, how it controls protein expression of its targets. Several previous reports have suggested DAZL played important roles as an enhancer of mRNA translation in germ cell development (33-35). Because meiotic arrest was the prominent phenotype in *Dazl* conditional KO testes and genes encoding components of the synaptonemal complex were identified as DAZL targets, we investigated how expression of *Sycp1, Sycp2* and *Sycp3* was affected by *Dazl* KO at translational level via polysome analysis. RNA was isolated from the fractions of free ribonucleoprotein (RNP) and polysomes (see Figs 5A and 5B). *Sycp1, Sycp2* and *Sycp3* transcripts were not significantly decreased in total transcripts but were dramatically decreased in polysome fractions (see Figs 4E and 5C). Hence, reduction of protein expression of those DAZL targets resulted from reduced protein translation rather than reduced mRNA level, further supporting a role for DAZL in translational regulation.

**Figure 5.**
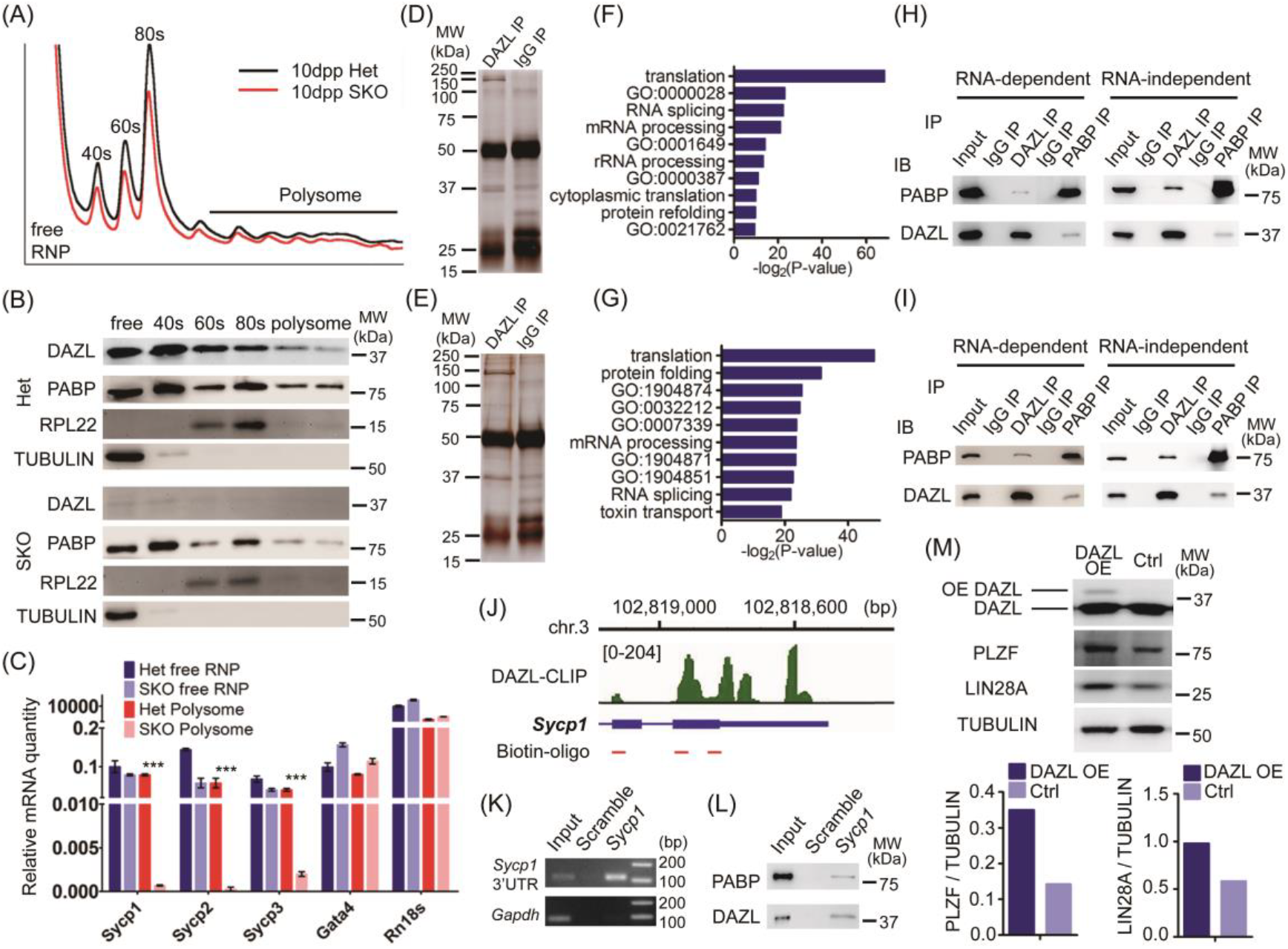
DAZL recruits PABP to regulate translation of its target. (A) Sucrose density gradient fractionation of 10dpp mouse testes with and without DAZL. (B) Immunoblot analysis of different fractions. (C) Target mRNAs from polysome fractions decreased dramatically with *Dazl* KO. (D) Silver staining of SDS-PAGE gel of DAZL and IgG immunoprecipitation (IP) in 8dpp mouse testes. Proteins were identified by mass spectrometry from excised bands (the whole lane except heavy and light chain). (E) Silver staining of SDS-PAGE gel of DAZL and IgG IP in 25dpp mouse testes. Proteins were identified by mass spectrometry from excised bands (the whole lane except heavy and light chain). (F) Gene ontology analysis results of proteins identified by mass spectrometry from (D). GO:0000028: ribosomal small subunit assembly; GO:0001649: osteoblast differentiation; GO:0000387: spliceosomal snRNP assembly; GO:0021762: substantial nigra development. (G) Gene ontology analysis results of proteins identified by mass spectrometry from (E). GO:1904874: positive regulation of telomerase RNA localization to Cajal body; GO:0032212: positive regulation of telomere maintenance via telomerase; GO:1904871: positive regulation of protein localization to Cajal body; GO:1904851: positive regulation of establishment of protein localization to telomere. (H) Co-IP of DAZL and PABP in 8dpp mouse testes. (I) Co-IP of DAZL and PABP in 25dpp mouse testes. (J) Biotin-oligo was designed to reverse complement the 3′UTR of *Sycp1* mRNA. (K) 3′UTR of *Sycp1* mRNA was enriched in *Sycp1* pull-downed samples. (L) DAZL and PABP proteins were enriched in *Sycp1* pull-downed samples. (M) Overexpression (OE) of DAZL in SSCs enhanced the expression of PLZF and LIN28A. Relative expression of PLZF and LIN28A in different samples was quantified using ImageJ.

Next, we performed mass spectrometry to identify proteins that interact with DAZL protein via IP (see Figs 5D and 5E). We identified more than 100 proteins that potentially interact with DAZL (see Supplementary Table S3). Among them, proteins involved in translation were significantly enriched, specifically, a series of ribosomal proteins and poly(A) binding protein (PABP), which stimulates translation when bound to a poly(A) tract (see Figs 5F and 5G, and Supplementary Figs S7A and S7B). PABP was previously found to interact with DAZL in frog oocytes (24). To determine if DAZL directly interacts with mouse PABP in the testis, we performed co-IP of DAZL and PABP in 8dpp and 25dpp mouse testis extract. DAZL and PABP reciprocally pulled down each other independent of RNA, establishing a direct interaction between these two proteins *in vivo* (see Figs 5H and 5I). We then ask if we could identify DAZL-PABP complex bound to a DAZL mRNA target directly by pulling down 3′UTR of the DAZL’s target. Using three Biotin-labeled Oligo probes complementary to the 3′UTR of a meiotic target of DAZL, *Sycp1*, we could pull down *Sycp1* 3′UTR from the testis extract (see Figs 5J and 5K). Remarkably, we could also pull down DAZL and PABP proteins at the same time (see Fig. 5L), demonstrating DAZL-PABP interaction directly on a DAZL target. To confirm the nature of translational regulation of DAZL, we next overexpress DAZL in SSC *in vitro*. The protein level of both DAZL targets *Plzf* and *Lin28a* was enhanced with DAZL overexpression (OE) (see Fig. 5M). Thus, DAZL function by recruiting PABP to mRNA targets of DAZL to promote their translation. Since DAZL bound to the 3′UTR of mRNA targets associated with SSC, meiosis and round spermatids, we propose that DAZL facilitates translation by recruiting PABP to those mRNA targets to form active translational circles (see Fig. 6). Without DAZL, translation of key proteins central for SSC maintenance, the synaptonemal complex, DNA repair and spermatid differentiation were severely impaired, resulting in a progressive loss of germ cells, failure of synapsis, arrest in meiosis and round spermatids. Indeed, we found that DAZL and PABP are concomitantly expressed from the SSC to round spermatid stages during spermatogenesis (see Supplementary Fig. S7C), and co-localize in spermatogenic cells in 10dpp and 25dpp mouse testes (white arrow, see Supplementary Fig. S7D). These findings established a central and global role of DAZL-mediated translational control throughout spermatogenesis (see Fig. 6).

**Figure 6.**
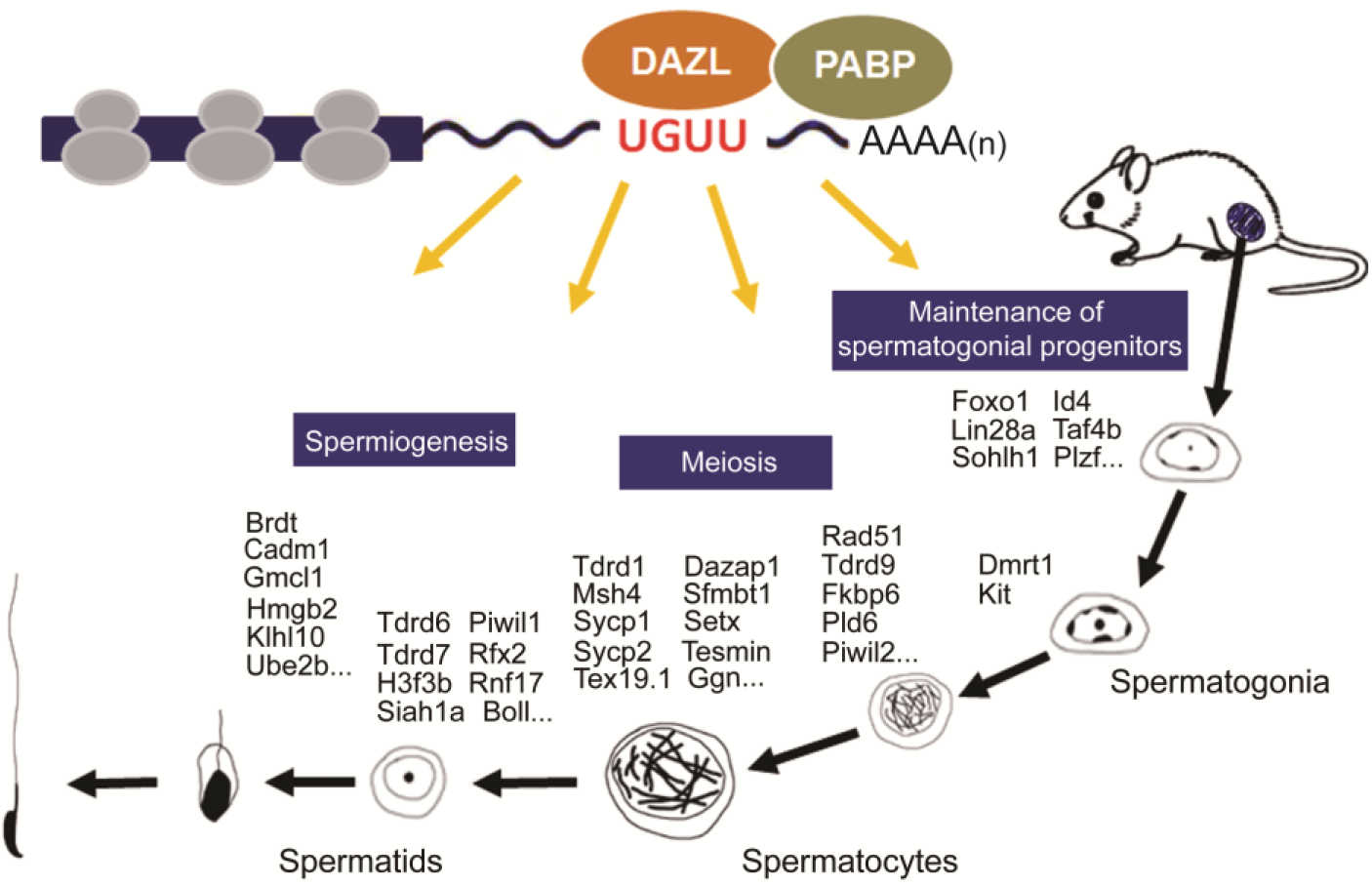
Model for DAZL-mediated master translational control in at least three key steps of spermatogenesis.

## DISCUSSION

Our finding suggests that DAZL expression is not only a hallmark of mouse germ cells, but is critically required throughout male gametogenesis but not in female gametogenesis. While earlier studies demonstrated a requirement for DAZL in fetal germ cell development and differentiation (20), our findings extended DAZL’s requirement into postnatal male gametogenesis and argued for a central regulatory role of DAZL throughout germ cell development, from embryonic to postnatal sperm development. Our genome-wide identification of DAZL targets and its protein partners in combination with systemic interrogation of DAZL requirement led us to propose that DAZL mediated translational regulation is critical throughout spermatogenesis and DAZL acts as a master translational regulator to ensure proper translation of key spermatogenic factors and thus fertility.

Ample genetic evidence on human DAZ family (DAZ, DAZL and BOULE) genes and their homologs among animals support their essential roles in human and animal fertility (8-11, 51, 52). However, molecular mechanisms by which DAZ proteins regulate sperm development remains elusive, with their direct *in vivo* targets largely unknown. Identification of more than three thousand direct targets from our transcriptome-wide HITS-CLIP experiments hence provided a key mechanistic piece for DAZL function. We showed DAZL protein directly regulate translation of their target mRNAs via binding to the 3′UTR of target mRNAs, without apparent effect on mRNA level, supporting a conserved translational function in mammalian germ cells (22-24, 33, 34, 53). Among a number of protein partners of DAZL identified through mass spectrometry, we showed that mouse DAZL interacts with PABP in the testis to promote translation of spermatogenic targets, extending the original finding in frog oocytes to their physiological context in the mouse testis (23, 24). Bioinformatics analysis of target sequences revealed 3mer GUU and 4mer UGUU being consensus binding motifs, consistent with the results from crystal structure analysis of DAZL RRM (30). Spermatogenic transcripts are highly enriched among DAZL targets, pathway analysis revealed potential roles in spermatogonial stem cell development, meiosis and spermatid differentiation, among which only two transcripts were previously reported to be regulated (33, 34).

Previously *Dazl* knockout germ cells exhibited a defect in spermatogonial transition from A_align_ to A_1_ type spermatogonia (17), we showed clearly a requirement of DAZL in spermatogonial stem cell maintenance. Conditional deletion of *Dazl* by either *Vasa*-cre or *Stra8*-cre led to a gradual loss of germ cells and SSCs marked by PLZF^+^ and LIN28A^+^ cells, with no apparent increase in KIT^+^ spermatogonia (unpublished). This finding established a role of DAZL in SSC maintenance but not in spermatogonia differentiation (54). The meiotic roles of DAZL were suggested as *Dazl* knockout germ cells fail to enter meiosis or were arrested at or before leptotene during the first wave of spermatogenesis (18, 55), we found that spermatogenic cells lacking DAZL apparently could differentiate and enter meiosis but were arrested in zygotene/pachytene transition stage, supporting critical meiotic requirement of Dazl (18). Difference in the specific arrest stage may reflect unique properties of knockout germ cells from the whole body DAZL knockout or limitation of very few germ cells available for analysis (18, 55).

DAZL’s role in spermatogonial stem cells revealed a novel layer of translational regulation in SSCs maintenance and differentiation. SSCs regulation involves a hierarchy of transcriptional regulators and signaling pathways (46, 56). The only posttranscriptional regulator required in spermatogonial stem cell maintenance is *Nanos2*, another RNA binding protein, (57), though the molecular mechanism by which NANOS2 regulates spermatogonial stem cell was not determined. Interestingly NANOS2 repressed DAZL expression in sexually differentiating germ cells at embryonic stages (58), it raised an interesting possibility that NANOS2 might exert its influence in SSC via DAZL. Indeed *Nanos2* overexpression resulted in a reduction of SSC chain and an increase of A_s_, similar to our *Dazl* knockdown phenotype (see Figs 3I and Supplementary Fig. S5C) and to *Dazl* conditional knockout effect on the testis (57). Hence NANOS2 may regulate the level of DAZL expression to balance the SSC maintenance and differentiation. Future experiments are needed to determine the contribution of NANOS2 and DAZL to SSC maintenance and their interaction in maintaining the homeostasis of spermatogenesis.

Master developmental regulators are the few key regulators in the developmental pathways, which regulate a large number of downstream targets in critical developmental steps. DAZL binds to many spermatogenic target transcripts and appear to regulate their translation to ensure sufficient protein expression for key steps of spermatogenesis. Hence DAZL may represent a master translational regulator to facilitate efficient sperm production from embryonic germ cell stage to postnatal spermatogenesis. Indeed DAZL is most extensively expressed from embryonic germ cell throughout postnatal spermatogenesis validating its expression as a hallmark of germ cells; Secondly, DAZL is required persistently throughout germ line development including most stages of spermatogenesis, without which male germ cell development is arrested thereafter (11, 19, 20) (This work); Thirdly, DAZL binds to a large number of spermatogenic transcripts key to the progression of spermatogenesis and recruits PABP to the 3′ UTR of those target transcripts. Without DAZL, translation of those key spermatogenic regulators is disrupted, resulting in an arrest in subsequent steps. Hence we proposed that DAZL function as a master translational regulator during spermatogenesis (see Fig. 6). The insights and mechanistic frameworks established by this work could advance our understanding not only of the fundamental mechanisms distinguishing germ cells from somatic cells but also of human infertility involving DAZ family proteins.

Added note: During the preparation of this manuscript, another manuscript reported their CLIP analysis on DAZL and their validation using the FACS sorted germ cells from whole body *Dazl* knockout testes (59). Two thirds of our targets overlapped with their targets, confirming the validity of our targets. However, Zagore *et al* found that a portion of target transcripts were down-regulated in the *Dazl* knockout spermatogenic cells, leading to their conclusion that DAZL regulated the stability of a subset of its target transcript. While our analysis showed that DAZL knockout does not affect the transcript steady level of DAZL target but mainly affect their translation. Such a difference may result from the different animal model used for RNAseq study as their *Dazl* knockout mice are full-body knockout while our model are stage-specific knockout model. Despite this difference, there is still a large portion of DAZL target transcripts level not changed in absence of DAZL from Zagore *et al*. This major portion of target transcripts unchanged in *Dazl* KO germ cells may correspond to the transcripts under translational regulation by DAZL from our study.

## METHODS

### Animals and cells

The *Dazl*^f/f^ ES cells used for this research project was generated by the trans-NIH Knock-Out Mouse Project (KOMP) and obtained from the KOMP Repository (www.komp.org). *Stra8*-cre, *Vasa*-cre and *Hspa2*-cre mice were obtained from the Jackson Laboratory. Wild type C57/B6 mice were purchased from the Animal Core Facility of Nanjing Medical University. All animal experiments were conducted with the guidelines approved by the Institutional Animal Care and Use Committee (IACUC) of Nanjing Medical University.

ESCs were derived from the inner cell mass (ICM) of mouse pre-implantation blastocyst embryos and maintained with N2B27/2iLIF medium supplied with 1% knockout serum replacement (Thermo Fisher Scientific, 10828028) on cell culture plates coated with 0.2% (w/v) gelatine. Primary SSCs were sorted by anti-THY-1 antibody conjugated-beads from 6-8 dpp mouse testes. Pachytene spermatocytes, round spermatids and elongated spermatids were purified by using STA-PUT apparatus. Long-term culture of SSCs was established following a previously described protocol (60).

### RNA isolation, qPCR and RNA-sequencing

Total RNA was extracted with TRIzol reagent (Invitrogen, 15596018) from the cells or whole testes. All reagents for RNA experiments were performed using nuclease-free water (Invitrogen, AM9937). The appropriate RNA was reverse-transcribed into cDNA with random primers (Takara, RR036A). qPCR was performed using a SYBR Green Master Mix Kit (Vazyme, Q141). Relative gene expression was analyzed based on the 2^−ΔΔCt^ method with *Gapdh* as internal control. At least three independent experiments were analyzed. All primers were listed in the supplementary Table S4.

For RNA-sequencing, isolated total RNA was converted into cDNA for RNA sequencing using Illumina Truseq RNA Sample Preparation Kit, followed by rRNA removal and sequenced on an Illumina HiSeq 2500 using 2 × 150nt sequencing.

### Western blot

We prepared protein samples by RIPA lysis buffer (50mM Tris (pH 7.4), 150mM NaCl, 1% NP-40, 0.5% sodium deoxycholate) containing a protease inhibitor cocktail (Roche, 11697498001). The primary antibodies used were as follows: rabbit anti-DAZL antibody (Abcam, ab34139, 1:1000); rabbit anti-α-TUBULIN antibody (Santa Cruz, sc-8035, 1:5000); rabbit anti-DDX4 antibody (Abcam, ab13840, 1:800); rabbit anti-SYCP1 antibody (Abcam, ab175191, 1:1,000); mouse anti-OCT4 antibody (Santa Cruz, sc-5279, 1:1000); goat anti-PLZF antibody (R&D, AF2944, 1:500); rabbit anti-LIN28A antibody (Abcam, ab46020, 1:2000); rabbit anti-STRA8 antibody (Abcam, ab49602, 1:1000); mouse anti-SPO11 antibody (Santa Cruz, sc-377161, 1:1000); rabbit anti-SYCP2 antibody (a gift of Dr. P.J. Wang, University of Pennsylvania, 1:500); mouse anti-SYCP3 antibody (Abcam, ab97672, 1:1000); rabbit anti-RAD51 antibody (Abcam, ab133534, 1:1000); rabbit anti-SMC3 antibody (Abcam, ab9263, 1:1000); rabbit anti-PUM2 antibody (Abcam, ab10361, 1:1000); rabbit anti-PABP antibody (Abcam, ab21060, 1:1000); rabbit anti-TNP2 antibody (a gift of Dr. Stephen Kistler at University of South Carolina, 1:1000).

### Histology, Immunofluorescence and immunohistochemistry

Paraffin-embedded testes were fixed with Hartman’s Fixative (Sigma, H0290) overnight at room temperature before and serially sectioned (5μm) to be stained with hematoxylin and eosin for histologic analysis. The sections were also used to perform immunofluorescence and immunohistochemistry analysis.

For immunostaining using frozen sections, testes were embedded in optimum cutting temperature compound (O.C.T.) and frozen in liquid nitrogen. Serial sections were cut at 5μM thickness onto glass slides and stored at −80°C until use.

The primary antibodies used were as follows: rabbit anti-DAZL antibody (Abcam, ab34139, 1:200); mouse anti-DAZL antibody (AbD Serotec, MCA2336, 1:50); rabbit anti-LIN28A antibody (Abcam, ab46020, 1:1000); mouse anti-H2AFX antibody (Millipore, 16-202A, 1:200); rabbit anti-MIWI antibody (Abcam, ab12337, 1:100); rabbit anti-DDX4 antibody (Abcam, ab13840, 1:200); rabbit anti-SYCP1 antibody (Abcam, ab15090, 1:100); rabbit anti-SYCP3 antibody (Abcam, ab15093 1:100); mouse anti-SYCP3 antibody (Abcam, ab97672, 1:100); rabbit anti-LIN28A antibody (Abcam, ab46020, 1:1000); goat anti-PLZF antibody (R&D, AF2944, 1:100); rabbit anti-PABP antibody (Abcam, ab21060, 1:200); rat anti-TRA98 antibody (Abcam, ab82527, 1:200); rabbit anti-SYCP2 antibody (a gift of Dr. P. Jeremy Wang, University of Pennsylvania, 1:200); rabbit anti-FOXO1 antibody (Cell Signaling Technology, 2880, 1:100); rabbit anti-TNP2 antibody (a gift of Dr. Stephen Kistler at University of South Carolina, 1:200); rabbit anti-Sohlh1 antibody (a gift of Dr. Aleksandar Rajkovic at the University of Pittsburgh, 1:200).

### Whole mount staining

Remove testes tunica and spread the tubules apart gently, then wash 3 times with 1 × Dulbecoo’s PBS and disperse the seminiferous tubules with 0.5mg/mL collagenase and 7mg/mL DNase I at 37°C for 2-10 mins depending on the testes. After washing with DPBS, the tubules were fixed in 4% PFA overnight at 4°C. To washing tubules 2 times with DPBS, remove DPBS and add 25% MeOH in PBST (0.1% Tween20 in PBS). To remove 25% MeOH and add 50% MeOH in PBST to the tubules, followed by incubate for 5 min on ice with gently rocking. Then removing the tubules to 75% and 100% MeOH step by step and storing at −20°C for at least 2h. Next to remove the tubules from 100% MeOH to 25% MeOH gradually and permeablized in PBS with 0.2% Triton X-100 and 0.9% H_2_O_2_ at room temperature for 40-50min. After washing in PBST for 5min, the tubules were blocked in 2% BSA with 0.1% Triton X-100 and 5% FBS for 1h at room temperature, and incubated in primary and secondary antibody, washed 3 times in PBST and mounted in Fluoroshield with DAPI (Sigma, F6057). The primary antibodies used were as follows: mouse anti-DAZL antibody (MCA2336, AbD Serotec, 1:50); rabbit anti-LIN28A antibody (ab46020, Abcam, 1:1000).

### HITS-CLIP

DAZL HITS-CLIP was performed essentially as for MOV10L1 previously (43). Testes from 25dpp mice were collected, de-tunicated, disrupted by mild pipetting in ice-cold HBSS, and immediately UV-irradiated three times at 254 nm (400 mJ/cm^2^). The cells were pelleted and washed with PBS, and the final cell pellet was flash-frozen in liquid nitrogen and kept at −80°C. UV light-treated cells were lysed in 300μL of 1× PMPG with protease inhibitors, 2U/μL RNasin inhibitor (Promega, N2515), and no exogenous nucleases; lysates were treated with DNase (Promega, M6101) for 5min at 37°C and then centrifuged at 90,000g for 30min at 4°C. For each immunoprecipitation, 10μg of anti-DAZL mouse monoclonal IgG was bound on protein A Dynabeads (Invitrogen, 10002D) in antibody-binding buffer (0.1M Sodium phosphate at pH8, 0.1% NP-40) for 3h at 4°C; antibody-bound beads were washed three times with 1× PMPG (1x PBS [no Mg^2+^ and no Ca^2+^], 2% Empigen). Antibody beads were incubated with lysates (supernatant of 90,000g) for 3h at 4°C. Low- and high-salt washes of immunoprecipitation beads were performed with 1× and 5× PMPG (5× PBS, 2% Empigen). RNA linkers (RL3 and RL5) as well as 3′ adaptor labeling and ligation to CIP (calf intestinal phosphatase)-treated RNA CLIP tags were previously described (61). Immunoprecipitation beads were eluted for 12 min at 70°C using 30μL of 2×Novex reducing loading buffer. Samples were analyzed by NuPAGE (4%–12% gradient precast gels run with MOPS buffer). Cross-linked RNA–protein complexes were transferred onto nitrocellulose (Invitrogen, LC2001), and the membrane was exposed to film for 1–2h. Membrane fragments containing the main radioactive signal and fragments up to ~5 kDa higher were cut (see Fig. 2B). RNA extraction, 5’ linker ligation, RT–PCR, and the second PCR step were performed with the DNA primers (DP3 and DP5 or DSFP3 and DSFP5) as described previously (61). cDNA from two PCR steps was resolved on and extracted from 3% Metaphor 1 × TAE gels stained with Gel-red. The size profiles of cDNA libraries prepared from the main radioactive signal and higher molecular weights were similar (see Fig. 2B). DNA was extracted with QIAquick gel extraction kit and submitted for deep sequencing. The cDNA libraries were sequenced on an Illumina HiSeq 2500.

### Peak calling for CLIP-seq

Homer software (http://homer.salk.edu/homer/ngs/index.html) was used to call peaks of CLIP-Seq data, when input file (RNA-Seq data) was given as background signal to remove noises. Tag directories for samples were constructed using makeTagDirectory in Homer package and peak regions were called using findPeaks in Homer package, too. The parameters for find Peaks are: -F 2 -L 2 -poisson 0.05 –style factor (fold enrichment over input tag count is set to 2, and fold enrichment over local tag count is set to 2, Set poisson p-value cutoff is 0.05);

### Motif analysis for called peak regions

A motif discovery algorithm (find Motifs Genome.pl) was designed for regulatory element analysis in Homer. It takes two sets of sequences and tries to identify the regulatory elements that are specifically enriched in one set relative to the other. It uses ZOOPS scoring (zero or one occurrence per sequence) coupled with the hypergeometric enrichment calculations (or binomial) to determine motif enrichment. Several motif lengths, default=3, 6, were considered for peaks regions found in each CLIP-seq library. The significantly enriched motifs for each library were reported and sorted by p-values. De novo predicted motifs were performed by findMotifsGenome method.

### RNA immunoprecipitation

Mouse testes were lysed in polysome lysis buffer (PLB, 0.5% NP40, 100mM KCl, 5mM MgCl**2**, 10mM HEPES, 1mM DTT, 100 units/mL RNase Out, 400μM VRC, and 1× protease inhibitor cocktail, pH7.0) by mechanical homogenizer and centrifuged at 20,000g for 20 min at 4°C to remove the debris. The supernatant was pre-cleared with blocked Dynabeads and then immunoprecipitated with antibody-conjugated or normal IgG-conjugated Dynabeads at 4°C for at least 3h. Next, the beads were washed four times with NT2 buffer (0.05% NP40, 50mM Tris-HCl, 150mM NaCl, and 1mM MgCl_2_, pH 7.4) supplemented with RNase and proteinase inhibitor. Before the last washing, the beads were divided into two portions. A small portion was used to isolate protein for enrichment identification of target protein, the remaining portion was resuspended in 100μL of NT2 buffer supplemented with RNase inhibitor and 30μg of proteinase K to release the RNA at 55°C for 30min, the RNA was eluted using 1 mL of TRIzol.

### Surface nuclei spread

Mouse testes were removed, seminiferous tubules gently minced with tweezers in DMEM, and cells mechanically separated. The cellular suspension was then spun to pellet cellular debris, and the nuclear suspension was pipetted onto slides. Slides were then fixed for 3 minutes each in freshly prepared 2% paraformaldehyde in PBS containing 0.03% SDS, and in 2% PFA alone. Slides were rinsed three times for 1 minute each in 0.4% PHOTO-FLO 200 solution (Eastman Kodak Company, Rochester, NY), dried, and then blocked in TBST containing 10% goat serum. Slides were then incubated with primary antibodies for 1 hour at 37°C or overnight at 4°C.

Primary antibodies used for immunofluorescence were as follows: rabbit anti-SYCP1 antibody (Abcam, ab15090, 1:100); rabbit anti-SYCP3 antibody (Abcam, ab15093 1:100); mouse anti-SYCP3 antibody (Abcam, ab97672, 1:100); mouse anti-H2AFX antibody (Millipore, 16-202A, 1:200); human anti-Crest antibody (15-234-0001, Antibodies inc, 1:100).

### Sucrose Density Gradient Fractionation

Dissected testes from 10dpp mice were homogenized in MBC buffer. After removal of nuclei and major organelles such as mitochondrion by centrifugation, supernatants were loaded onto continuous 20–50% sucrose gradients prepared in lysis buffer without detergent. Gradients were centrifuged for 3h at 180,000g in a SW41 rotor (Beckman, Mountain View, CA). Fractions (0.5 ml) were collected manually on BIOCOMP gradient machine (Canada), and the A_260_ was measured. RNA was isolated from 20% of gradient fractions with TRIzol. Five percent of each fraction was immunoblotted with DAZL antibody. For developmental polysome analysis, testes from 10dpp wild type and *SKO* mice were prepared and fractionated as above.

### Co-immunoprecipitation

Mouse testes were lysed in Pierce IP Lysis Buffer (Thermo Scientific, 87787) containing complete EDTA-free protease inhibitor cocktail (Roche, 04693132001), then immunoprecipitated with anti-DAZL antibody (AbD Serotec, MCA2336) and anti-PABP antibody (Abcam, ab21060) in the presence or absence of 5U/μL RNase A and 200U/μL RNase T1 (Thermo Scientific, AM2286) by Pierce Crosslink IP Kit (Thermo Scientific, 26147). The co-immunoprecipitated proteins complexes were detected using western blotting with the following antibodies: rabbit anti-DAZL antibody (Abcam, ab34139, 1:1000) and rabbit anti-PABP antibody (Abcam, ab21060, 1:1000).

### Plasmid construction

For Dazl expression, plasmid pCDH-Dazl, pCMV6-mouse Dazl (mDazl-WT) and pCMV6-mouse Dazl RRM deletion (mDazl-del) were generated by PCR amplification of the coding sequences (CDS) followed by recombining into pCDH-EF1-MCS-T2A-Puro (SBI, CD520A-1) or pCMV6-Entry destination plasmid (Origene, PS100001) using the ClonExpress MultiS One Step Cloning Kit (Vazyme, C113). For knockdown experiments, shRNA-coding DNA fragments were synthesized and subcloned into the AgeI and EcoRI sites of the pLKO.1 TRC-cloning vector (a gift of Dr. Chen Dahua, State Key Laboratory of Reproductive Biology, Beijing, China) to create pLKO.1-shDazl plasmid (sh285 and sh2383). For Dual-Luciferase Reporter (DLR) Assay, various Dazl targets sequences including Dazl binding region were amplified by PCR and inserted into psiCHECK-2 destination plasmid (Promega, C8021). Mutant sites were generated by synthesized primers and recombined into psiCHECK-2 using the ClonExpress MultiS One Step Cloning Kit (Vazyme, C113). All primers were listed in the supplementary Table S4.

### Dual-Luciferase Reporter (DLR) Assay

When the 293T cells were 70–90% confluent, wild type and RRM deletion Dazl expression plasmids were transfected with the psiCHECK-2 plasmid which containing Dazl binding region into the cells simultaneously using Lipofectamine LTX Reagent (Promega, 15338). The cells were collected 48h after transfection and manipulated according to the manufacturer’s instructions of Dual-Luciferase Reporter Assay System (Promega, E1910).

### *Dazl* shRNA Knockdown

For generation of the lentivirus to knockdown Dazl expression in SSC, 293T cells were transfected with a combination of pLKO.1-shDazl plasmid and packaging plasmids using pmd-REV, pmd-1G/pmd-LG (a gift of Dr. Chen Dahua, State Key Laboratory of Reproductive Biology, Beijing, China). A pLKO.1 empty vector was used as negative control. Lentivirus-containing supernatant was collected 48h after transfection. After centrifugation at 1500g for 10min and filtration by 0.45μM filter membrane, the supernatant was centrifuged by ultracentrifugation at 50000g for 2h at 4°C. The deposition was resuspended in HBSS, and then stored at −80°C for future use.

### Mass spectrometry

DAZL protein complex pull-downed by anti-DAZL antibody from mice testes at 8dpp and 25dpp were analyzed by mass spectrometry Core Center, Nanjing Medical University using the entire elutes.

### Biotin-Oligo pulldown of RNA

Adult mouse testes were digested to single cells and cross-linked with 1% formaldehyde in PBS for 10 min at room temperature, and cross-linking was then quenched with 0.125 M glycine for 20 min at 4°C. The cells were lysed in RIPA buffer (150mM NaCl, 50mM Tris-HCl, 5mM EDTA, 1% NP-40, 0.1% SDS) with freshly added 1mM DTT, protease inhibitors, and 0.1 U/μl RNase inhibitor) for 30 min at 4°C followed by sonication and centrifugation. Biotin-labeled Oligos (100pmol), complementary to 3′ UTR of Sycp1 mRNA, were added to the supernatant, which was mixed by end-to-end rotation at 37°C for 4h. M-280 Streptavidin Dynabeads (Life Technologies) were washed three times in RIPA buffer, which was blocked with 500 ng/μl yeast total RNA and 1mg/ml BSA for 2h at room temperature, then washed three times again in RIPA buffer before being re-suspended. 100μl washed/blocked Dynabeads was added per 100pmol of biotin-DNA oligos, and the whole mix was then rotated overnight at 4°C. Beads were captured and washed five times with 40× the volume of Dynabeads with RIPA buffer. Beads were then subjected to RNA and protein elution.

### Data availability

Data for RNA sequencing of 10dpp testes from control and *Dazl*^f/-^;*Stra8*-Cre males have been deposited in the Gene Expression Omnibus database under accession code GSE114042. Data for Dazl-CLIP of 25dpp testes from wild type C57/B6 mice have been deposited in the Gene Expression Omnibus database under accession code GSE113710. The authors declare that all data supporting the findings of this study are available within the article and its supplementary information files or from the corresponding author on reasonable request.

## Author contributions

E.Y.X. conceptualized the research; E.Y.X. and H. L. designed the study; H.L. performed most of the experiments; Z.L., J. Y., D.W., H.W., J.C. and E. Y. X. performed and interpreted experiments; B.G. performed bioinformatics analyses for DAZL-CLIP; E.Y.X. supervised the study and wrote the paper with H.L.

## ACKNOWLEDGEMENTS

The *Daz*^f/f^ ES cells used for this research project were generated by the trans-NIH Knock-Out Mouse Project (KOMP) and obtained from the KOMP Repository (www.komp.org) at UC Davis and CHORI (U42RR024244). We would like to thank Drs. Xin Wu and Ke Zheng for advice and assistance on HITS-CLIP experiments and SSCs experiments respectively, as well as discussion throughout the project. We thank Jieli Chen, Guihua Du, Fan Yang, Xinrui Wang and Jiachen Sun for technical assistance, Dr. Renee Reijo Pera and Youqiang Su for comments on the manuscript. We thank Genergy Biotech, Shanghai, China for bioinformatics analysis. We thank Dr. P. Jeremy Wang for the SYCP2 antibody, Dr. Stephen Kistler for the TNP2 antibody and Dr. Aleksandar Rajkovic for the SOHLH1 antibody.

This work was supported by grants from National Basic Research Program of China (973 program, 2015CB943002 and 2013CB945201), National Natural Science Foundation of China (31771652, 81270737 and 81401256), and Natural Science Foundation of Jiangsu Province (BK2012838).

---

**Fig. S1.**
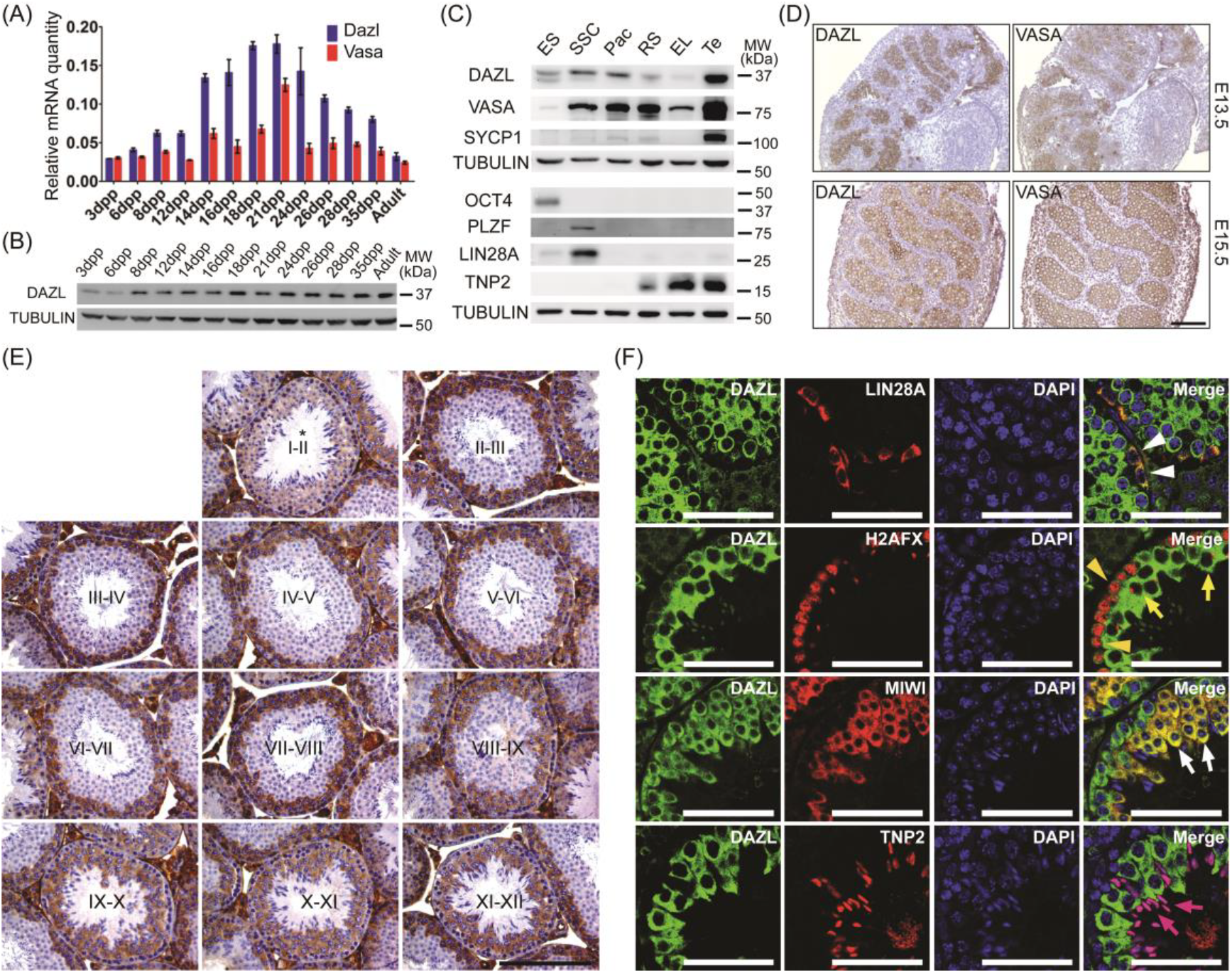
*Dazl* dynamically expressed throughout male germline development, from embryonic gonad, SSCs to round spermatids. (A) Realtime PCR of *Dazl* RNA in mouse testes at different day post-partum (dpp) during the first wave of spermatogenesis. (B) Western blot analysis of DAZL protein in mouse testes at different dpp during the first wave of spermatogenesis. (C) Western blot analysis of DAZL protein in various purified or cultured cells. ES, Embryonic stem cells; SSCs, Spermatogonial stem cells; Pac, Pachytene spermatocytes; RS, Round spermatids; EL, Elongating spermatids; Te, Testes. (D) DAZL and VASA staining in E13.5 and E15.5 mouse testis sections. Scale bars: 100μm. (E) DAZL staining in mouse testes sections at different stages of testicular epithelial cycles. Scale bars: 100μm. (F) Dynamic stage expression of DAZL in mouse testes with stage-specific markers. SSCs (white arrowhead); leptotene/zygotene (yellow arrowhead); pachytene (yellow arrow); round spermatid (white arrow); elongated spermatid (violet arrow). Scale bar: 50μm.

**Fig. S2.**
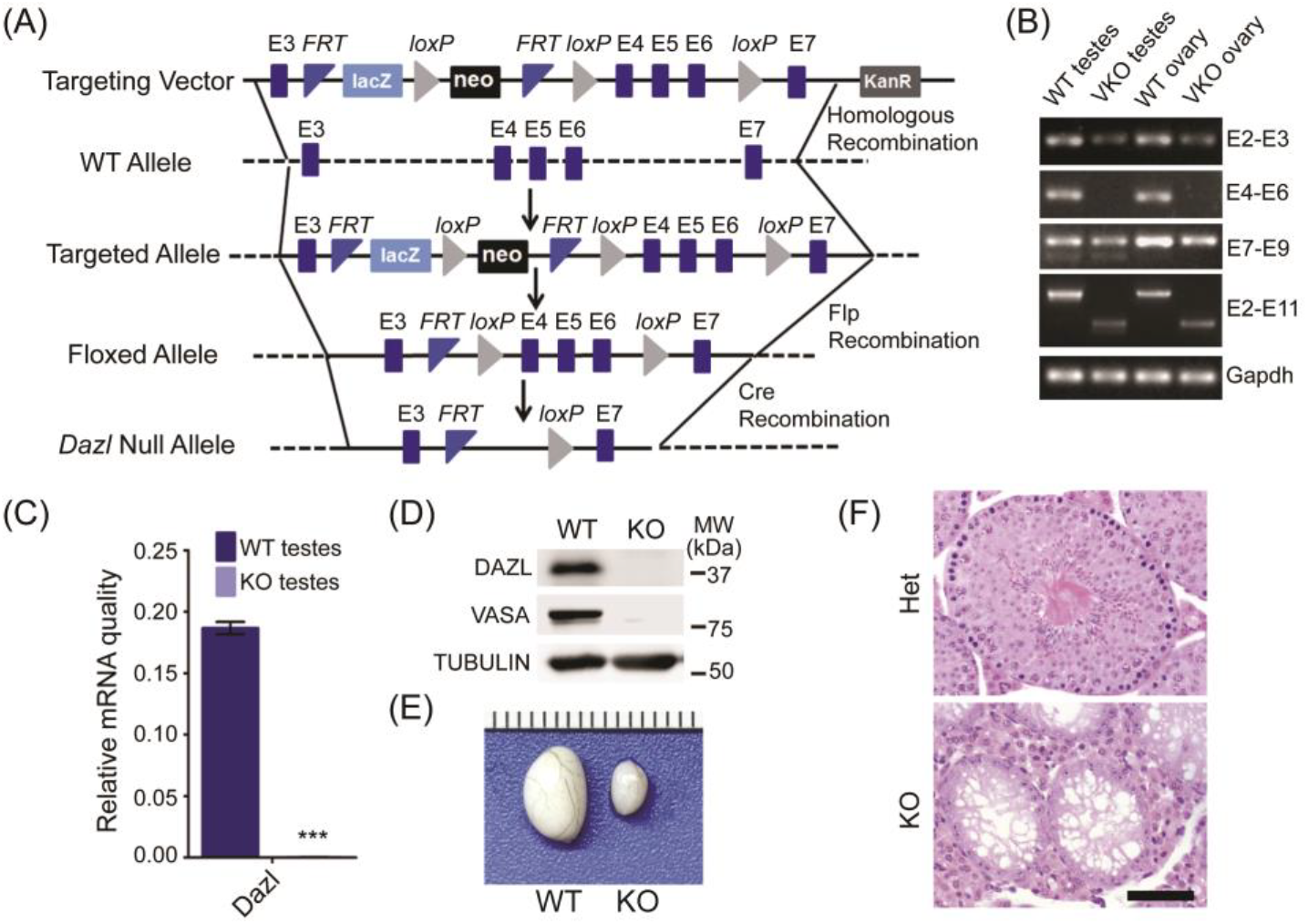
Generation of *Dazl* conditional knockout mice. (A) Targeting strategy of the *Dazl* gene. (B) WT transcripts containing exon 4-6 (E4-E6) were absent in *Dazl* VKO tissues. (C) qRT-PCR analysis of *Dazl* transcripts between exon 4 to exon 6 in *Dazl* VKO testes, as normalized to *Gapdh*. (D) DAZL and VASA were absent in adult knockout testis due to lack of germ cells. (E) Testes excised from adult *Dazl^Δ^/^Δ^* mice were very small. (F) Histological defects in adult het and *Dazl^Δ^/^Δ^* mice testis. Scale bars: 200μm.

**Fig. S3.**
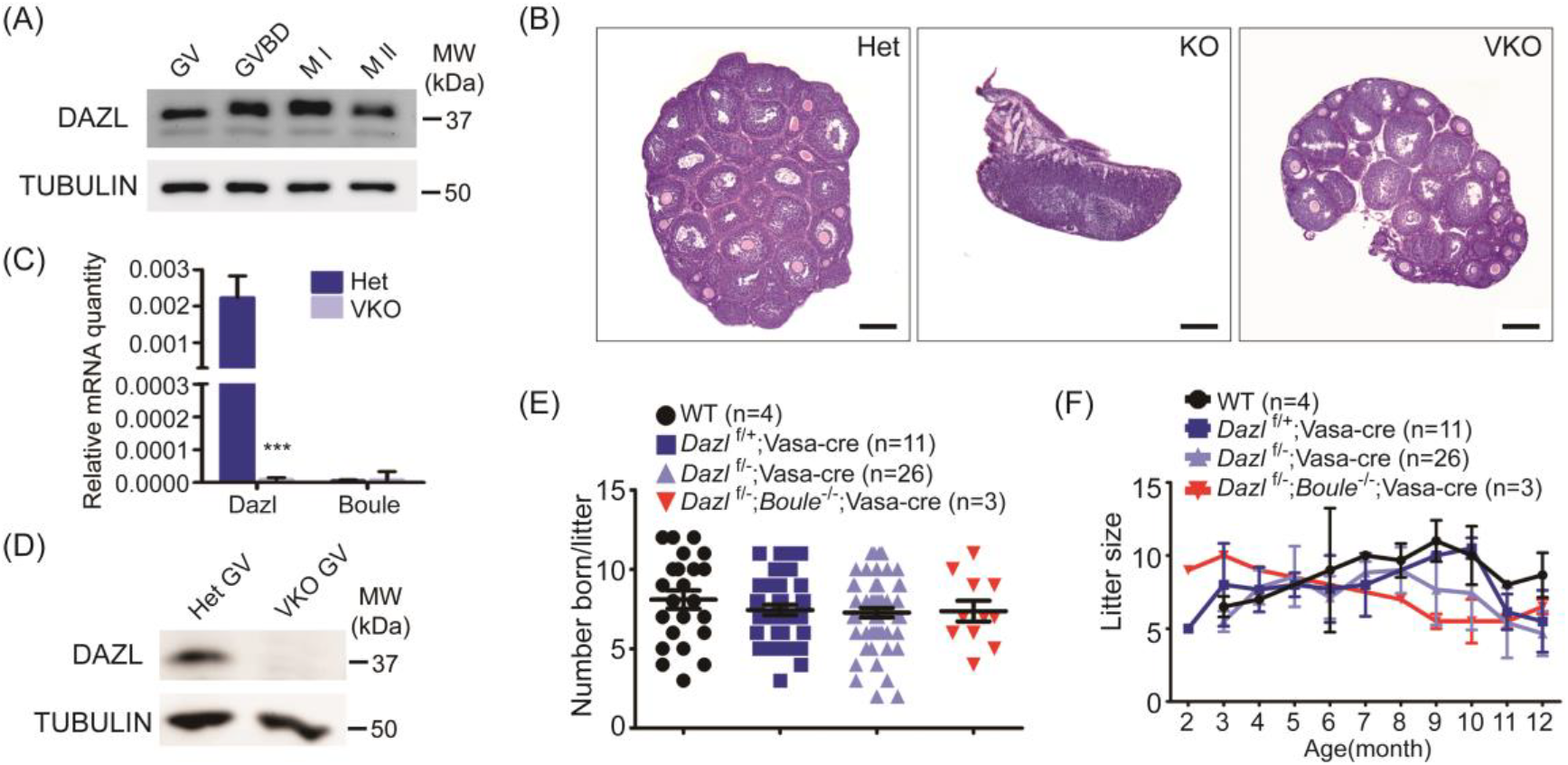
*Dazl* is not required for female fertility. (A) DAZL exists throughout oogenesis. (B) H&E stained ovary sections from 21dpp Het, *Dazl* KO and *Dazl* VKO mice showing *Dazl* is required for oogenesis but not folliculogenesis. Scale bar: 200μm. (C) qRT-PCR analysis of *Dazl* transcripts between exon 4 to exon 6 in the ovaries, as normalized to *Gapdh*. (D) DAZL was absent in *Dazl* VKO GV oocytes.. (E) Litter sizes of wildtype, heterozygotes, *Dazl* VKO and *Dazl Boule* double knockout females mated with a wildtype male continuously collected over 12 months. (F) Litter size distribution of wildtype, heterozygotes, *Dazl* VKO and *Dazl Boule* double knockout females with age. Error bars are SD. Using linear regression analysis.

**Fig. S4.**
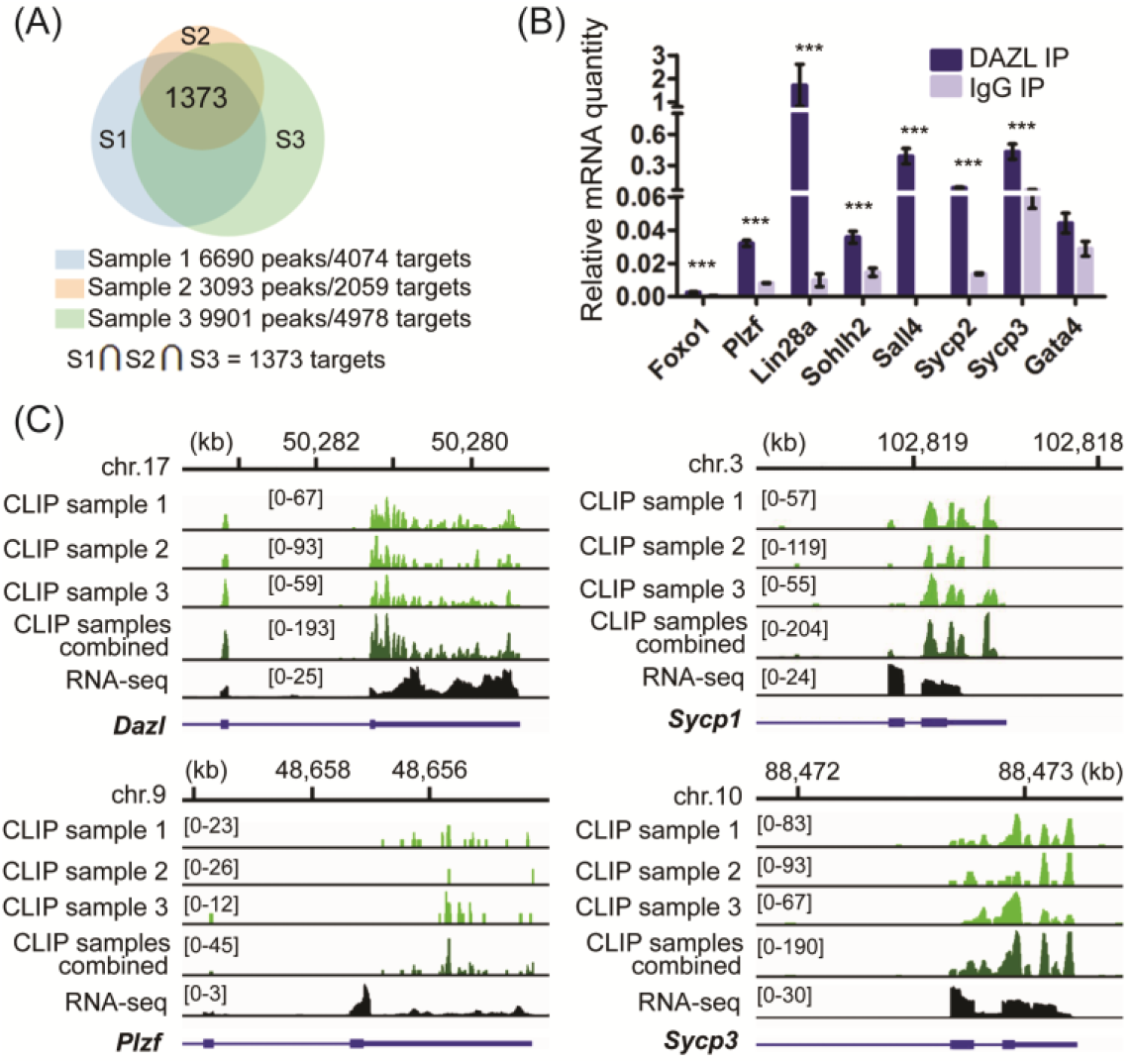
DAZL binding sites and motifs identified by CLIP. (A) Overlap of DAZL mRNA targets in two biological replicates, including samples from two main bands and a mixture of higher bands. (B) Validation of HITS-CLIP result by RIP on another seven target genes. (C) Schematic representation of highly correlated peak distribution from 3 samples and the combined peak representation will be used thereafter.

**Fig. S5.**
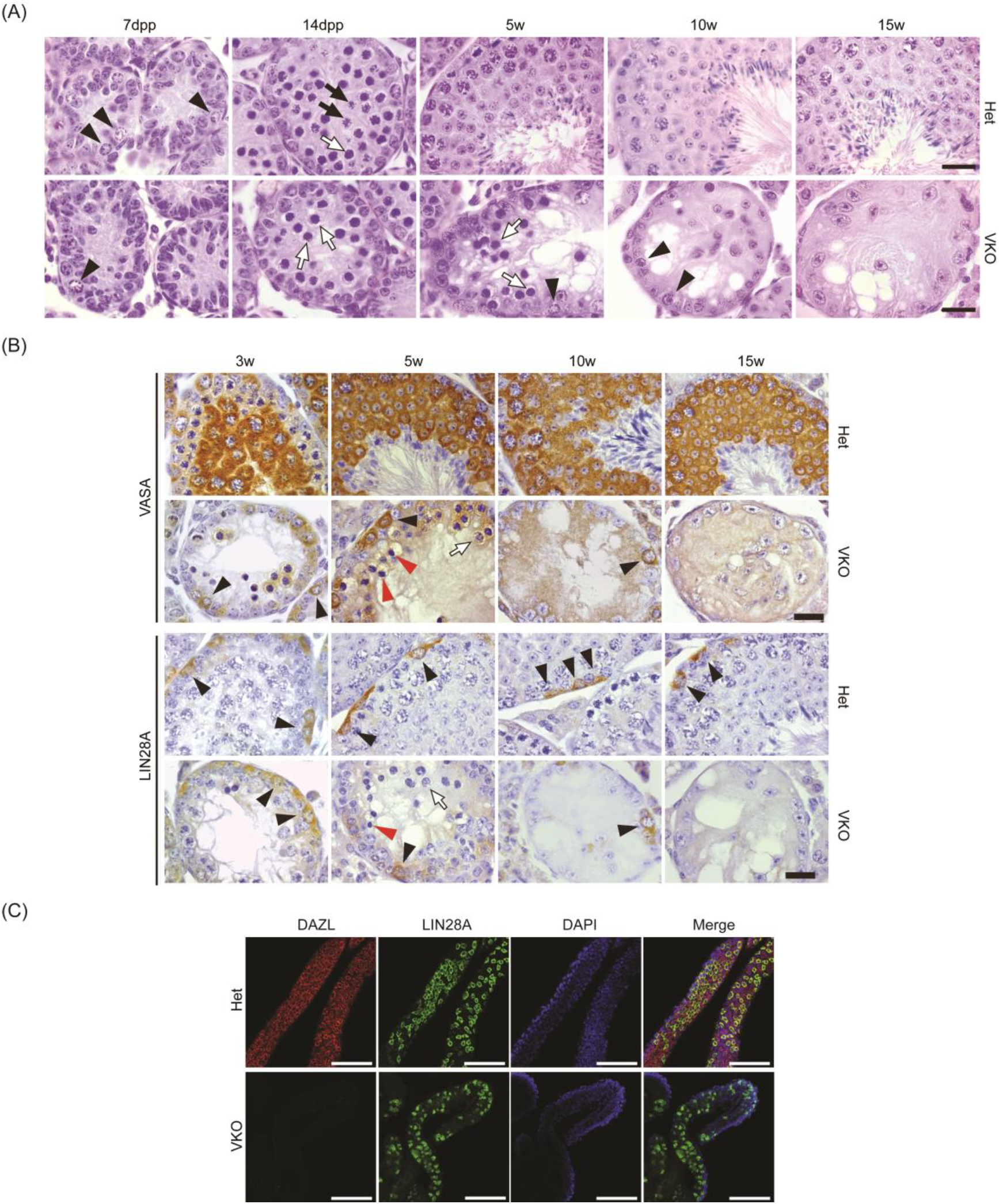
SSCs were lost gradually in *Dazl* VKO testes. (A) H&E stained testis sections from Het and VKO seminiferous tubules. Spermatogonia (black arrowhead), zygotene spermatocytes (white arrow) and pachytene spermatocytes (black arrow). (B) IHC staining of VASA and LIN28A in heterozygote and *Dazl* VKO testes. Scale bar: 20μm. (C) Whole mount staining of 10dpp heterozygote and *Dazl* VKO testes. Scale bar: 100μm.

**Fig. S6.**
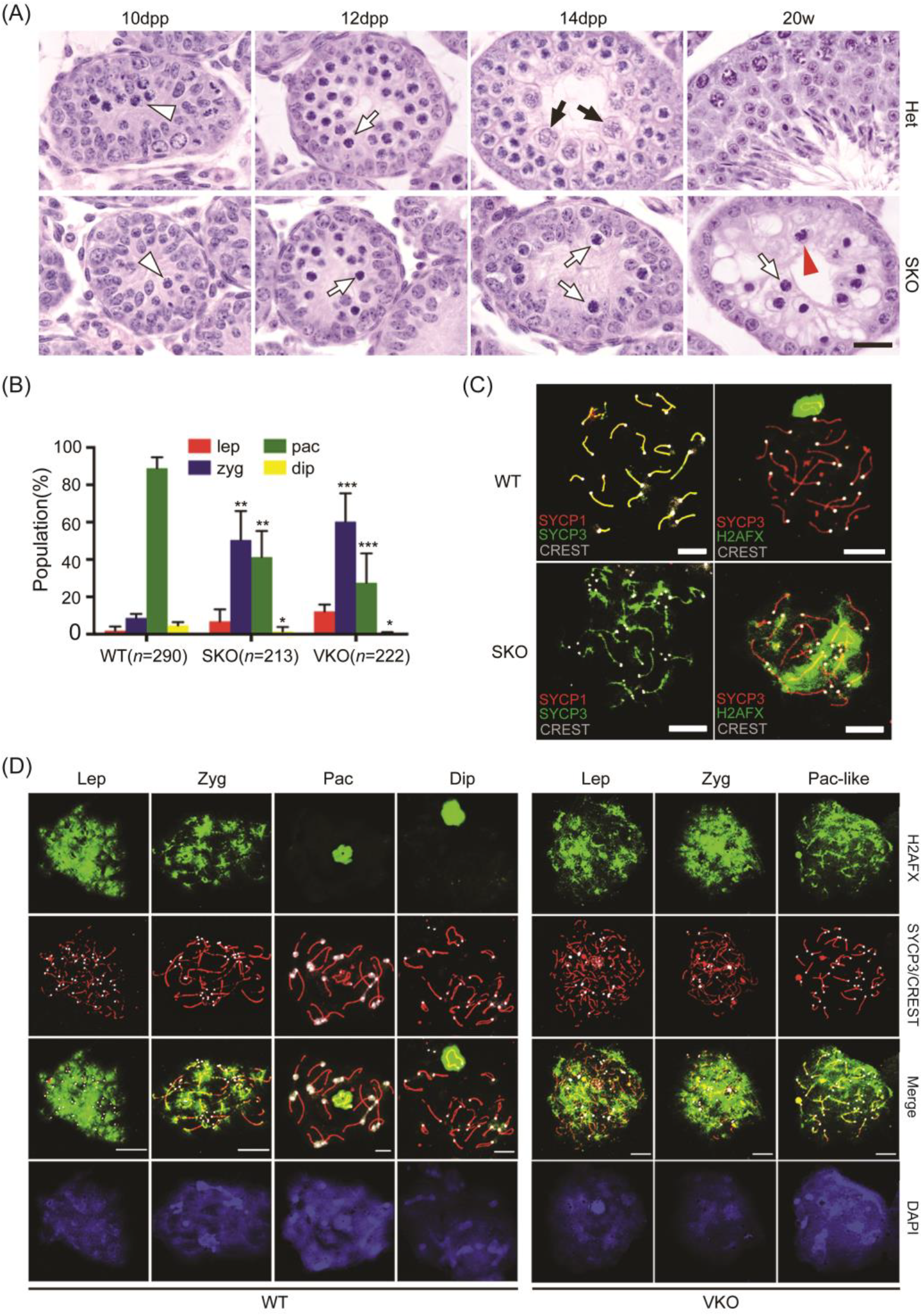
Accumulation of H2AFX in *Dazl* conditional knockout spermatocytes and meiotic arrest at zygotene-pachytene transition. (A) H&E stained testis sections from Het and SKO seminiferous tubules. Leptotene spermatocytes (white arrowhead), zygotene spermatocytes (white arrow), pachytene spermatocytes (black arrow) and typical apoptotic cells germ cells with dark condensed nuclei (red arrowhead). (B) Quantification of prophase I stages among meiotic cells based on the pycnotic chromosome morphology and degree of alignment of AEs (n: cells per genotype used). Error bars indicate SEM. *P<0.05, **P<0.01, ***P<0.005. Scale bars: 10μm. (C) Representative images of meiotic wild type and *Dazl* SKO spermatocytes co-stained with H2AFX, SYCP3 and CREST (white points) antibodies. In SKO mice, cloud-like nuclear staining of H2AFX remained at pachytene-like stage spermatocytes. For pahcytene-like spermatocytes, approximately 40 centromeres can be identified. Scale bars: 10μm. (D) Accumulation of H2AFX in pachytene and diplotene of 21dpp *Dazl* VKO mice. Representative images of meiotic wild type and *Dazl* VKO spermatocytes co-stained with H2AFX, SYCP3 and CREST (white points) antibodies. In *Dazl* VKO mice, cloud-like nuclear staining of H2AFX remained at pachytene-like stage spermatocytes. For pahcytene-like spermatocytes, approximately 40 centromeres can be identified. Scale bars: 10μm.

**Fig. S7.**
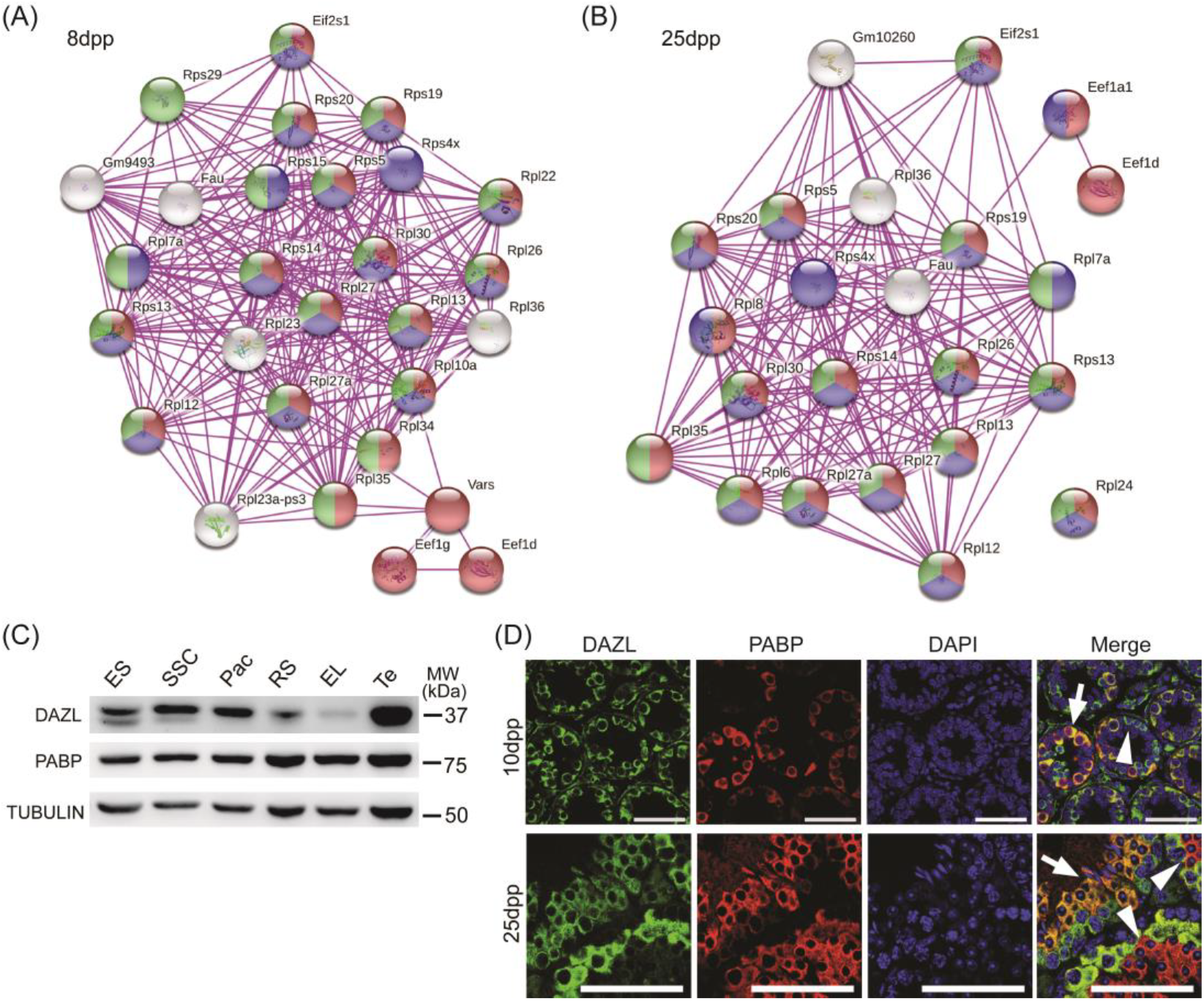
DAZL co-localized with PABP in germ cells and also interacted with many other proteins involved in translational control. (A) Western blot analysis of DAZL and PABP protein expression in male germ cells at different stages. (B) Co-localization of DAZL and PABP in mouse testes at 10dpp and 25dpp. Co-localized cells (white arrow), no co-localized cells (white arrowhead). Scale bar: 50μm. (C) DAZL interacting proteins involved in translational control at 8dpp testes. Pac, Pachytene spermatocytes; RS, Round spermatids; EL, Elongating spermatids; Te, Testes. (D) DAZL interacting proteins involved in translational control at 25dpp testes. Red ball indicated ‘translation’ gene ontology, blue ball indicated ‘poly(A) RNA binding’ gene ontology, green ball indicated ‘ribosome’ gene ontology. Purple lines indicate an interaction evidence from known experiments.

